# Individuals at risk for developing rheumatoid arthritis harbor differential intestinal bacteriophage communities with distinct metabolic potential

**DOI:** 10.1101/2021.02.03.429590

**Authors:** Mihnea R. Mangalea, David Paez-Espino, Kristopher Kieft, Anushila Chatterjee, Jennifer A. Seifert, Marie L. Feser, M. Kristen Demoruelle, Meagan E. Chriswell, Alexandra Sakatos, Karthik Anantharaman, Kevin D. Deane, Kristine A. Kuhn, V. Michael Holers, Breck A. Duerkop

## Abstract

Rheumatoid arthritis (RA) is an autoimmune disease characterized in seropositive individuals by the presence of anti-cyclic citrullinated protein (CCP) antibodies. RA is linked to the intestinal microbiota, yet the association of microbes with CCP serology and their contribution to RA is unclear. We describe intestinal phage communities of individuals at risk for developing RA, with or without anti-CCP antibodies, whose first degree relatives have been diagnosed with RA. We show that at-risk individuals harbor intestinal phage compositions that diverge based on CCP serology, are dominated by Lachnospiraceae phages, and originate from disparate ecosystems. These phages encode unique repertoires of auxiliary metabolic genes (AMGs) which associate with anti-CCP status, suggesting that these phages directly influence the metabolic and immunomodulatory capability of the microbiota. This work sets the stage for the use of phages as preclinical biomarkers and provides insight into a possible microbial-based causation of RA disease development.

## INTRODUCTION

Rheumatoid arthritis (RA) is a systemic autoimmune disease with a global prevalence of approximately 1%. The development of RA in at-risk individuals is dependent on a combination of genetics, epidemiological factors, and systemic immune dysregulation [1]. The heritability of RA is estimated to be 40–60%, with increased familial risk evident among first-degree relatives (FDRs) of individuals with diagnosed RA [2, 3]. Analyses of at-risk FDRs, even those without serum RA-related autoantibodies, have identified patterns of mucosal inflammation whereby anti-cyclic citrullinated peptide (anti-CCP) antibodies and rheumatoid factors (RF), as well as cytokines and chemokines, are expressed locally in a subset of individuals [4-6]. In addition, anti-CCP and RF are present in the blood for years prior to the onset of RA, and their presence as well as circulating cytokine and chemokine biomarkers, are predictive of future RA development [7-9]. To probe the mucosal origins hypothesis [1] and the mounting evidence implicating intestinal microbiota perturbations in RA etiopathogenesis [10], it is necessary to characterize the ecological associations of the microbiota in at-risk individuals susceptible to RA.

Studies linking the role of the intestinal microbiota to systemic autoimmune diseases predominantly rely on 16S ribosomal gene analyses of bacteria within the microbiome, and have expanded our understanding of dysbiosis in the RA intestine. Individuals with established RA harbor a microbiota dominated by *Prevotella copri* [11, 12], enriched with Gram-positive bacteria [13], and decreased carriage of bifidobacteria [14], Gram-negative *Bacteroides*, and Firmicutes [13, 15]. The association of enriched Prevotellaceae, including *P. copri*, has also been described in individuals with preclinical RA [16], indicating that intestinal *P. copri* is immune-relevant to the pathogenesis of RA [17]. The presence of *P. copri* may therefore represent a biological indicator and additional risk factor for RA development and progression [18]. However, associating a single organism to RA etiology neglects the interactions of bacteria with their surrounding environment and other bacterial community members whose populations can be influenced by predatory bacteriophages (phages).

In contrast to the recent enthusiasm for characterizing microbial links to the etiology of RA, relatively little is known concerning the composition of phage communities in the intestine as it relates to RA disease risk. Phages of the intestinal microbiota can fluctuate in community composition in response to immune system function and disease, which suggests that they could be exploited as biomarkers for early disease detection [19]. Metagenomic sequencing strategies have revealed extensive and diverse populations of phages in the human intestine [20-22], in which phage community dynamics correlate with distinct disease states [23-25]. Specific intestinal phage genomic signatures precede autoimmunity development of type 1 diabetes in a cohort of diabetes-susceptible children, with disease-associated phages correlating to the bacterial component of the microbiota [26]. In addition to the direct impact of intestinal phages on bacterial community composition via classical predation and prophage mediated bacterial competition and metabolism, phages also adhere to mucosal surfaces, significantly impacting microbial colonization [27] and host mucosal immunity development [28]. Evidence is emerging that phages are also immunomodulatory through intrinsic anti-inflammatory properties, and are capable of direct lymphocyte regulation through the ability to translocate to multiple tissues and organs [29]. Despite these observations and potential implications for systemic autoimmune diseases like RA, evaluation of intestinal phages in the context of RA disease risk has yet to be described.

The interplay between intestinal bacteria, their phages, and the host immune system, whose interactions have consequences not only for compositional dysbiosis but immunomodulation, must be considered in the etiopathogenesis of RA. The microbiome, and more recently the virome, have been implicated in a range of human diseases including cancers [30, 31], inflammatory bowel diseases [32, 33], and arthritis [11, 34]. By characterizing the phage populations in an at-risk RA FDR cohort; further sub-grouped with regard to autoantibody status as defined by the presence of anti-CCP antibodies and compared to healthy controls, we have begun to address this question. The cohort contains individuals that do not have inflammatory arthritis or established RA disease but are FDRs to an individual with diagnosed RA, which alone increases RA risk. Studying the microbiomes of at-risk individuals in the preclinical RA state could lead to the identification of biomarkers and therapeutic targets independent of confounding by the use of drugs in subjects with active arthritis.

We used metagenomics to define intestinal phage populations of anti-CCP positive (CCP+) and negative (CCP-) individuals in an at-risk FDR cohort. Phage matching to bacterial hosts showed divergent intestinal phage communities dependent on anti-CCP serology status. We observed an overabundance of phages targeting Bacteroidaceae and Sreptococcaceae bacteria in CCP+ at-risk FDRs as well as phages targeting Bacteroidaceae bacteria in CCP-at-risk FDRs. Importantly, analysis of the metabolic traits encoded in phage metagenomes revealed intra-cohort profiles reflecting distinct immunomodulatory potential. Phages with auxiliary metabolic genes (AMGs) that modify lipopolysaccharide and other outer membrane glycans of host bacteria were differentially abundant, implicating modifications to bacterial antigenicity [35] and bacterial fitness [36] in RA-associated communities. Core phage metabolic genes, including 14 genes which are globally conserved among phages from multiple diverse environments [37], as well as bacterial surface modifying enzymes, were associated with phages targeting *Flavonifractor* sp. in the CCP+ cohort and *Bacteroides* sp. in the CCP-cohort. Phages targeting Lachnospiraceae (*Clostridium scindens*) and Actinomyces (*A. oris*), including several AMGs, were over-abundant among CCP+ and CCP-individuals, respectively, compared to healthy controls. Our data show that there are unique and abundant intestinal phages specific to RA-susceptibility status, and this highlights their potential as biomarkers for preclinical RA and the need for further pursuit of community-level bacteria-phage interactions during the development and progression of RA.

## RESULTS

### First-degree relatives to individuals with rheumatoid arthritis

A total of 25 human subjects were identified from the Studies of the Etiology of Rheumatoid Arthritis (SERA) [38], including 16 FDRs of individuals with RA and 9 age and sex matched healthy controls (HC). FDR subjects for which a detectable level of anti-CCP autoantibody was present (defined by a value of ≥ 20 units/mL in either ELISA assay for anti-CCP3.1 IgA/IgG or anti-CCP3 IgG (Inova Diagnostics) [39]) were designated the CCP+ group (n = 8). FDRs with no anti-CCP detected were designated the CCP- group (n = 8) (Table 1). Mean ages for the three groups in this study were 61.3 ± 11.0 for CCP+, 49.0 ± 15.7 for CCP-, and 44.4 ± 13.6 for HC. The distribution of sexes for each group is reported as percent female, with 88.9% for CCP+, 62.5% for CCP-, and 66.7% for HC. Among the CCP+ and HC groups, 3/9 and 2/9 of individuals have reported ever smoking (a risk factor associated with RA), respectively (Table 1).

**Table 1.** Summary of the Subjects’ Characteristics for the Samples Included in the Study.

| VARIABLE | HC | CCP+ | CCP- |
| --- | --- | --- | --- |
| Count | 9 | 8 | 8 |
| Age (mean) | 44.4 | 61.3 | 49 |
| Age (SD) | 13.6 | 11 | 15.7 |
| Sex (% female) | 66.7 | 88.9 | 62.5 |
| Serum CCP+ (%) | 0 | 100 | 0 |
| Ever smokers (%) | 22.2 | 33.3 | 0 |

### Generation and curation of *de novo* assembled VLP contigs

We used individual fecal samples from the subjects obtained at the time of autoantibody and clinical evaluations, and isolated total genomic DNA for shotgun metagenomic sequencing using an untargeted amplification-independent approach [23, 40]. Samples were physically separated into whole metagenome (M), including all genomic DNA present in the sample, and virus-like particle (VLP) fractions, which were subjected to phage-specific precipitation (Figure S1A). Illumina sequencing resulted in an average of 123.8 ± 32.2, 135.2 ± 40.4, and 104.7 ± 45.9 million (M) paired end reads per sample for CCP+, CCP- and HC whole metagenomes, respectively, and an average of 67.3 ± 29.5, 73.2 ± 33.7, and 89.6 ± 47.8 M paired reads per sample for CCP+, CCP- and HC VLP fractions, respectively (Figure S1B). VLP sequencing reads were used for *de novo* contig assembly of VLP metagenomes. In total, 3.56 M contigs were assembled and pooled from the 25 individual metagenomes, with 80,762 contigs longer than 5 kb (Figure 1A). VLP contigs longer than 5 kb were distributed evenly across the three sample groups, totaling 2908.6 ± 1461.3, 3209.0 ± 2573.8, and 3535.7 ± 2826.4 contigs per sample for CCP+, CCP- and HC respectively (Figure S1C).

**Figure 1.**
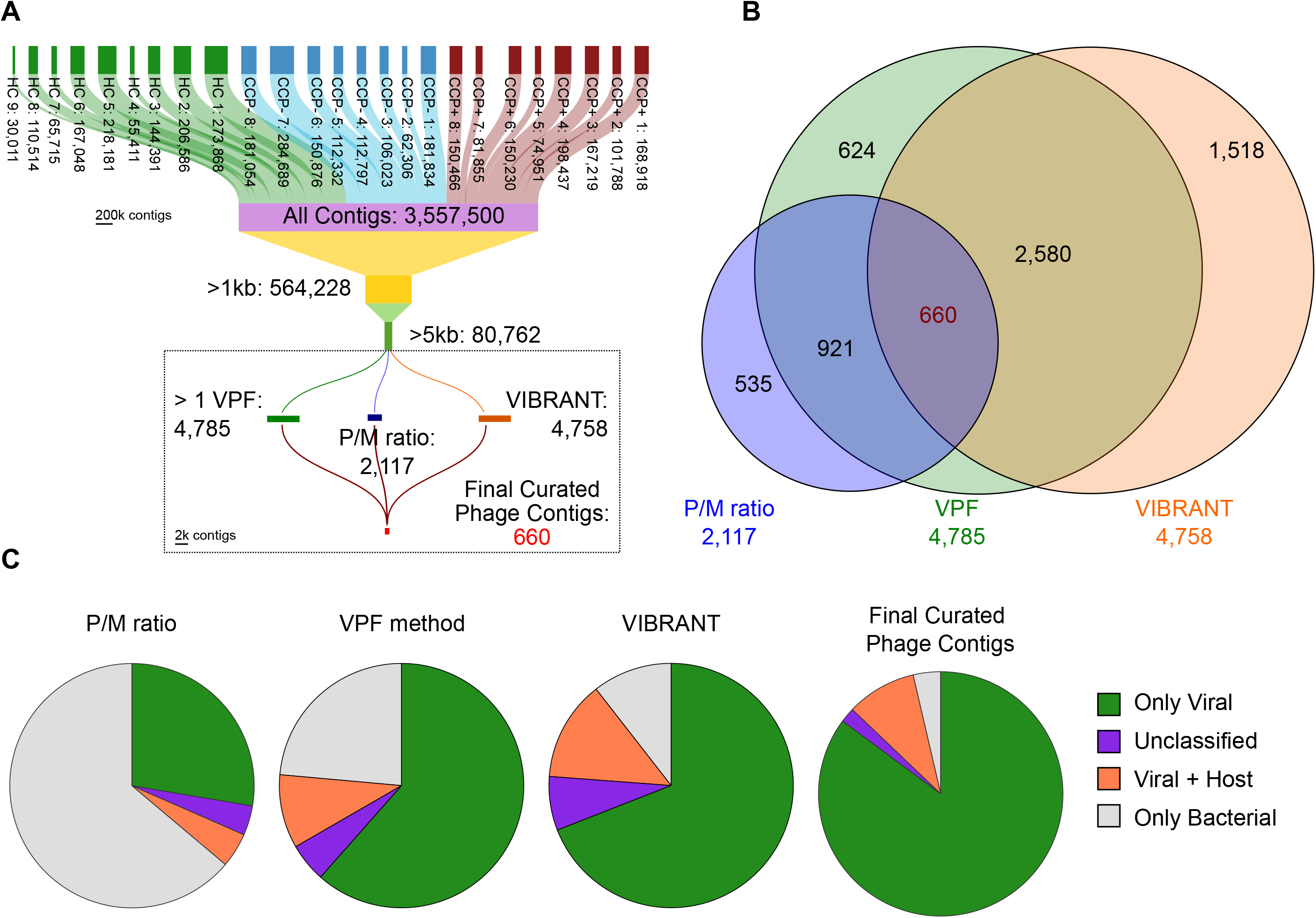
Generation and curation of *de novo* assembled VLP contigs. Metagenomic sequencing was carried out for 25 samples belonging to 3 cohorts of individuals, FDRs at risk for developing RA later in life with either CCP+ or CCP-serology status, and a Healthy Control (HC) group. (A) Contigs were assembled *de novo* for all samples, ranging from 30,011 to 284,689 contigs per sample, and a total of 3,557,500 contigs for the entire sample set. Each node on the Sankey diagram is scaled to the number of contigs it contains. Thresholds of minimum contig sizes being greater than 1 and 5 kilobases reduced the total numbers to 564,228 and 80,762 contigs respective to the size cut-off. Three independent methods were used to identify putative phages from the list of 80,762 contigs (boxed portion of panel A), resulting in 2,117 contigs from the P/M ratio method, 4,785 contigs from the Viral Protein Families method, and 4,758 contigs using the VIBRANT algorithm. (B) A Venn diagram was created to show the overlap of redundant contigs identified among the three methods. 660 unique contigs were identified independently by all phage identification methods. (C) CheckV analysis of the three separate methods as well as the final set of curated contigs revealed a disparity in host contamination, with the set of 660 contigs being relatively free of host bacterial contamination. Colors were assigned to the CheckV categories that account for prophage elements based on their position on the contig sequence, as well as pure viral (green) and pure bacterial (grey) classifications.

These 80,762 contigs served as a starting point for identifying putative phages using a three-pronged approach of independent phage discovery methods (Figures 1A and S1D). The first method (P/M ratio) employed a previously validated read mapping strategy whereby VLP read sets from all 25 samples were mapped to both whole metagenome (M) and VLP (P) contigs [23]. Using the read-mapping P/M ratio (see Methods), we identified 2,117 unique putative phage contigs after dereplication at 95% sequence identity. Next, we identified an independent set of phage contigs by aligning all open reading frames (ORFs) of the 80,762 VLP contigs against a set of 25,281 curated viral protein families (VPFs) [41]. Using this VPF method, several filters were applied to identify viral contigs; (i) 2,902 contigs were identified as having 5 or more VPF hits and non-viral Pfam hits below 20% of total ORFs on a contig, (ii) 263 contigs were identified with 5 or more VPF hits and less than 50% non-viral Pfam hits on a contig, (iii) 644 contigs with 2-4 VPF hits and 0 non-viral Pfams, (iv) 976 contigs with at least 1 VPF hit, without considering any non-viral Pfams. In total, after dereplication, the viral contigs arising from all above filters resulted in 4,785 unique viral contigs. For the third and final approach we employed VIBRANT (Virus Identification By iteRative ANnoTation), a sequence-independent algorithm that uses neural networks of viral protein signatures to identify lytic and lysogenic phages [37]. Using VIBRANT, we identified 4,758 unique viral contigs.

To consolidate this list, we identified contigs that were shared between all three phage discovery methods, resulting in a curated list of 660 contigs (Figures 1A and 1B). This curated list of putative phage contigs range in size from 5,007 bp to 557,525 bp. To assess host bacterial contamination among these contigs, we employed CheckV, a pipeline for assessing the quality of viral genomes [42]. CheckV analysis revealed a reduced level of host bacterial contamination and an increase of pure viral genomes in the final list of 660 curated contigs as compared to varying levels of contamination among the three separate methods prior to contig overlap identification (Figure 1C). We estimated completeness of our curated contigs using CheckV and determined a greater distribution of “high quality” contigs relative to contig length, in comparison to the three independent methods (Figure S2) [43]. Further, using the VIBRANT platform for integrated provirus prediction, we describe communities of predominantly lytic viral genomes belonging to Siphoviridae morphology (Figure S3). By using a combination of approaches for viral contig discovery and assessing the overlap among these methods, we have extracted a set of 660 predicted phages which are of overall high quality, both in terms of viral contig completeness and lack of bacterial contamination than those from each of the individual methods (Figures 1C and S2), which to date have been used primarily in isolation to identify and characterize viral metagenomes.

### Clustering of metagenomic viral contigs reveals distinct viral ecological composition

Next we compared our set of curated contigs to over 2.3 million viral whole genome and metagenome sequences from the IMG/VR database [44]. We used blastn at a threshold of 95% sequence identity over 85% of 1 kb sequence length and Markov clustering to group our contigs with related sequences from IMG/VR. Of the 660 contigs, 346 (52.4%) clustered into 255 clusters that contained 7,736 additional metagenomic viral contigs (mVCs) from IMG/VR. The remaining 314 contigs (47.6%) were classified as singletons, with an even distribution among CCP cohorts compared to healthy controls (Figure S4A). Of the curated contigs that were clustered, cluster sizes ranged from 2 to 646 members with 78.4% of the groups containing more than 2 partners and 36.5% containing more than 10 members, and 65.9% between 2 – 10 members (Figure S4B). Among these 255 clusters, 14 included reference prophages and lytic phages, and 318 (48.2%) clustered with classified mVCs, thus assigning multiple levels of taxonomy to our contigs (Figures 2A, 2B, and Supplementary Table 1).

**Figure 2.**
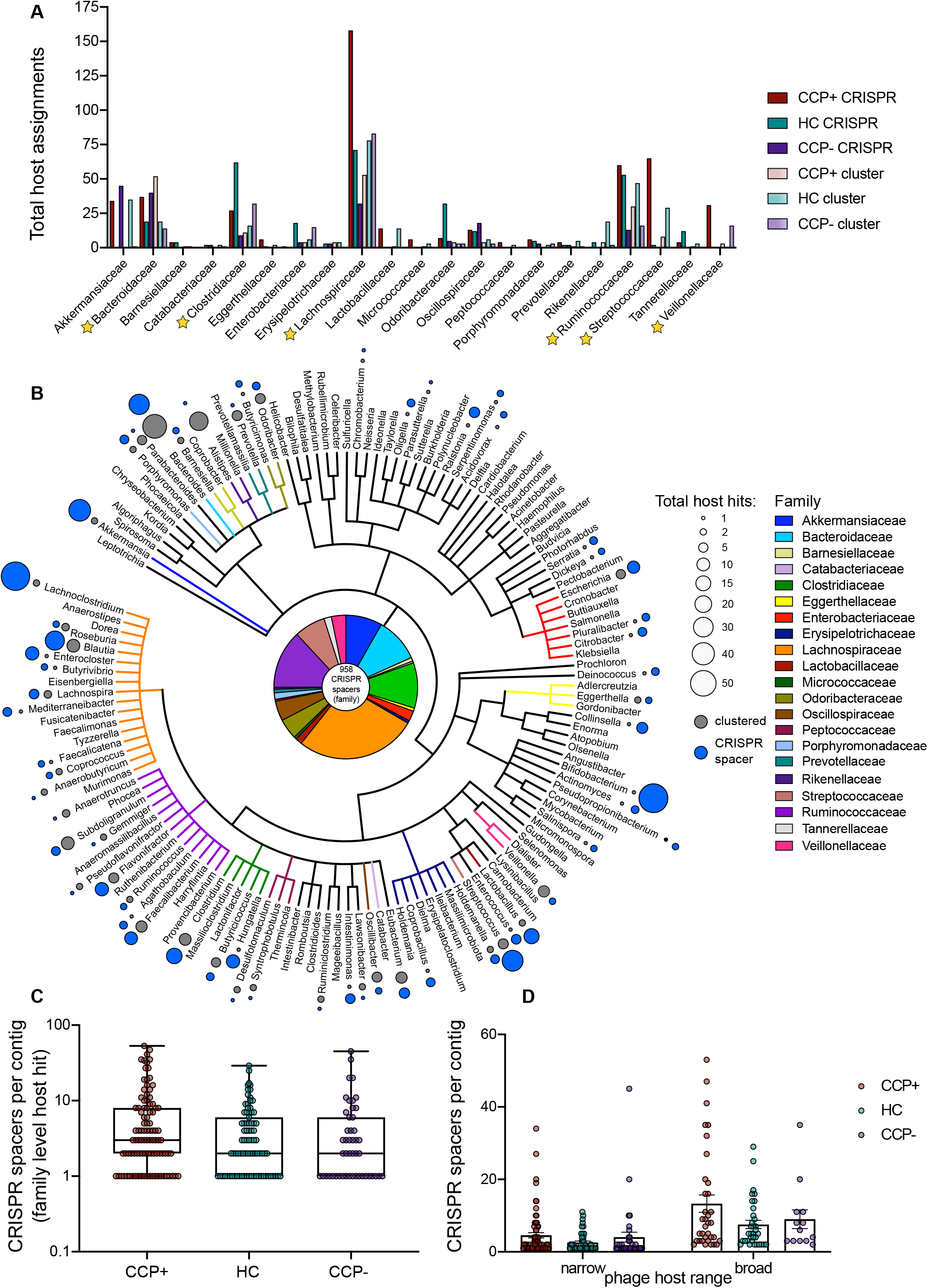
Clustering with metagenomic viral contigs reveals viral ecological composition. (A) Host assignments for the set of curated phages based on Markov clustering to the IMG/VR database metagenomic viral clusters or direct match to bacterial CRISPR spacers, based on cohort abundance. Bacteroidaceae, Lachnospiraceae, Ruminococcaceae, and Streptococcaceae hosts are evident to be cumulatively more abundant than other bacterial families, especially for the CCP+ cohort. (B) Cladogram of the complete host phylogeny at the genus level for all spacers identified from total RA virome via the VPF method. The pie chart at the center represents all 958 CRISPR spacers from the family level quantified in panel A that have been color coordinated on this cladogram as well. Total host hits were quantified at the genus level and are represented in relative size by colored circles, indicating host assignments that were discerned via clustering (dark grey) and those that were identified via direct CRISPR spacer matching (light grey). Total CRISPR spacers per contig with family level host taxonomy assignments were tabulated per cohort group (C) and differentiated as narrow or broad phage host ranges (D) based on target uniformity to bacterial family.

Although host assignments were made using sequence-based clustering, host specificity was further determined by aligning Clustered Regularly Interspaced Short Palindromic Repeat (CRISPR) spacer sequences to our 660 curated contigs. CRISPR-Cas serves as a snapshot of previous phage infections in the form of acquired spacer sequences that represent invading viral genomes [45], and these sequences can be used for accurate identification of phage-host interactions in intestinal microbiomes [23, 46]. CRISPR spacer host assignments at the family level were present in 207 of 660 contigs (31.4%). All CRISPR spacer queries considered for these analyses, ranging in length from 18 to 70 bp, were matches of 93.1–100% identity across the full length of the query and allowing for 0–2 mismatches and up to 1 gap throughout [47] (Supplementary Table 2). Among predicted phages, total assigned CRISPR spacers were evenly distributed, yet CCP+ sample containing phages predicted to target Lachnospiraceae, Ruminococcaceae, Streptococcaceae, and Veillonellaceae bacterial families were disproportionately abundant (Figures 2A and 2B). In total, 21 bacterial families were identified as hosts via CRISPR spacer matching, supplementing the phage-host interactions discerned from sequence-based clustering (Figure 2A). Among all samples in this study, phages were predicted to target Lachnospiraceae, Ruminococcaceae, Clostridiaceae and Bacteroidaceae bacteria with highest frequency of total CRISPR spacers (Figure 2A). Phage-host interactions were also measured in terms of host range specificity, showing that while the majority of the phages were predicted to have narrow host ranges, several spacers were linked to multiple hosts across family level and higher taxa (Figure 2C), consistent with prior observations of diverse viromes [47]. Broad host range phages were found across all cohorts, but particularly among CCP+ sample contigs (Figure 2D) suggesting a more dysbiotic community of host bacteria among these individuals’ metagenomes.

We further explored the association of sample cohorts to phage hosts using read mapping to determine differential host abundance profiles (Figure 3). Reads from all samples were mapped to assembled phage contigs whose host assignments were deduced using CRISPR-spacer matching and Markov clustering to quantify sequence abundances by measuring cohort-based read recruitment [23, 48-50]. In comparing the differential read recruitment to phages predicted to infect separate bacterial families, we observed differences based on reads originating from either the CCP+ or CCP-groups in relation to the HC cohort (Figure 3). Among the most striking, phage contigs targeting Bacteroidaceae recruited significantly more reads from CCP+ viromes than either HC or CCP-individuals (Figure 3A). In contrast, phages predicted to target Clostridiaceae bacteria were evenly abundant across all three groups (Figure 3B). For Lachnospiraceae bacteria, CCP+ phages recruited were evenly distributed among the groups with a slight elevation in CCP+ individuals that was not statistically significant (Figure 3C). Ruminococcaceae phages were significantly skewed when comparing HC to CCP-individuals (Figure 3D) and a major shift in phage read recruitment abundance was evident for Streptococcaceae phages, as a greater percentage of total CCP+ reads were mapped to these phages in relation to either HC or CCP-virome reads (Figure 3E). This skew among CCP+ individuals is supported by prior works showing elevated Streptococcal phage abundances in intestinal viromes of humans with inflammatory bowel disease [32] and a murine model of colitis [23]. Lastly, no significant differences were observed for read recruitment to Veillonellaceae-targeting phages (Figure 3F). Thus, differences in the host specificities were evident between CCP+, CCP-, and HC groups with respect to read mapping abundance profiles for Bacteroidaceae, Ruminococcaceae, and Streptococcaceae phages.

**Figure 3.**
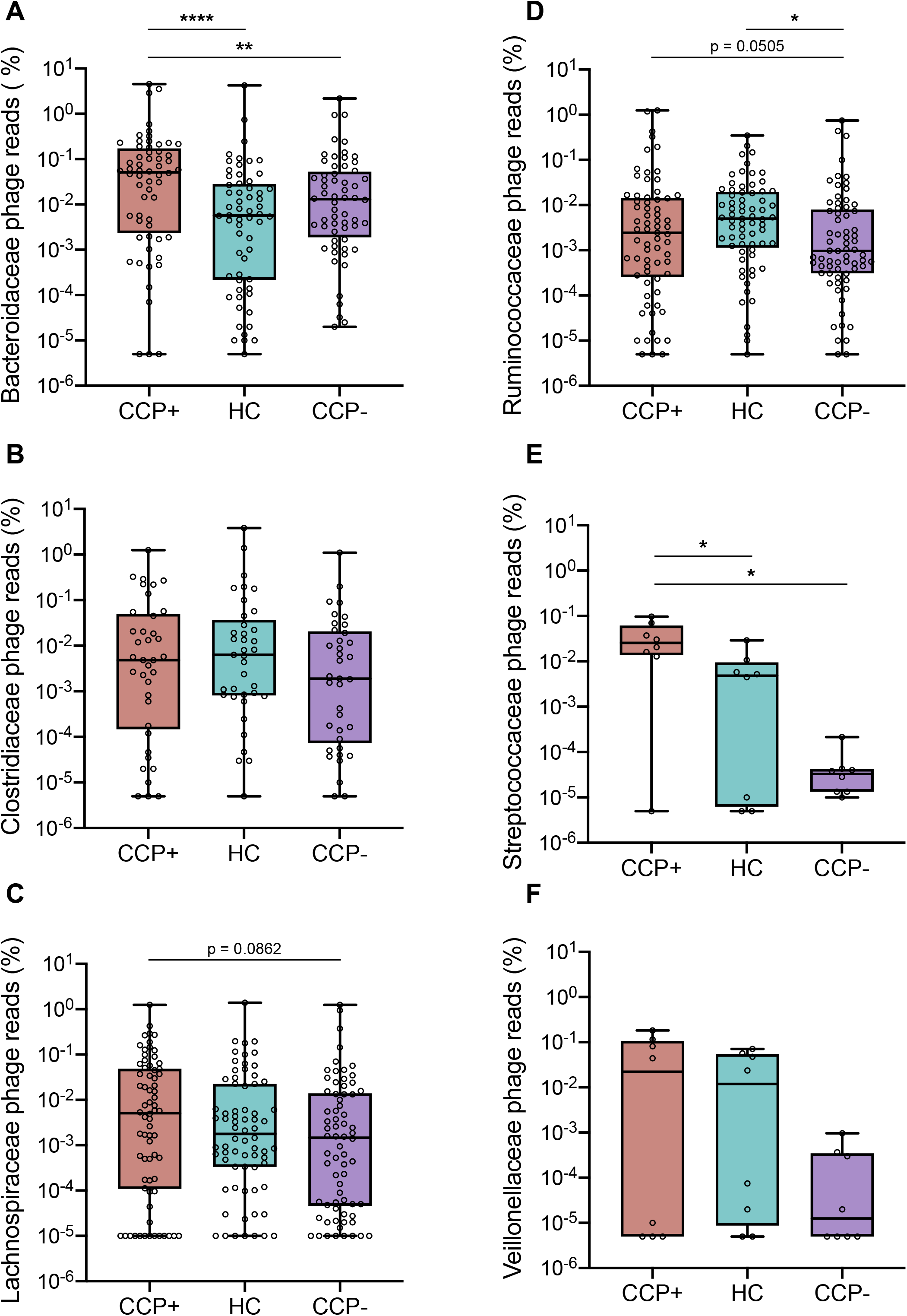
Phage-host assignments for curated VLP contigs reveal cohort-based differential read recruitment among several bacterial families. Relative abundances were calculated for all VLP reads mapped to phages predicted to target Bacteroidaceae (A), Clostridiaceae (B), Lachnospiraceae (C), Ruminococcaceae (D), Streptococcaceae (E), and Veillonellaceae (F) bacterial families. For these analyses, VLP reads were mapped to predicted phage contigs to which CRISPR spacers were assigned using bbmap at a 97% minimum read-mapping identity level. Scaffold abundances were averaged across all samples and statistics were determined by nonparametric Wilcoxon tests (* *p* < 0.05, ** *p* < 0.01, **** *p* < 0.0001).

### CRISPR spacer host metadata reveal CCP+ phages represent greater variability in microbial host ecology

To further explore the phage ecology from our subject cohort, we analyzed the host and mVC metadata from the Joint Genome Institute’s (JGI) Genomes OnLine Database (GOLD) [51]. The JGI GOLD database contains metadata from over 100,000 biosamples and over 350,000 sequencing projects involving genomic and metagenomic sequencing data from biological isolates worldwide. Moreover, recent work has contributed an additional 52,515 metagenome-assembled genomes from diverse ecologies and geographic distributions [52], further enhancing microbial host ecosystem analysis. Using the GOLD Biosample Ecosystem Classification system, we analyzed the ecosystem distributions for all CRISPR spacers identified in our curated contig list and discovered that the majority of host assigned contigs fell within four distinct ecosystem classification levels; from broad to specific environments: host-associated, human-associated, digestive system, and large intestine (Figure 4). For phages that were previously identified as having CRISPR spacer host assignments, total spacer alignments as identified by blastn ranging from 1 to 825 per contig, were tallied and used to calculate the uniformity of spacer origins per contig (Supplemental Table 3). For each of the four ecosystem categories, the most abundant classifications were used to compare across the study cohorts. At the highest order GOLD Ecosystem distribution, the host-associated (i.e., human, mammal, plant, arthropod, fungi) origin classification per contig was comparable for the HC and CCP-groups but not for the CCP+ group (Figure 4A). A similar pattern was evident at the lower order metadata distributions, with phage contigs derived from CCP+ individuals being more divergent from the other cohorts for contigs of human-associated origin (Figure 4B), digestive system origin (Figure 4C), and large intestine origin for the Ecosystem Subtype (Figure 4D).

**Figure 4.**
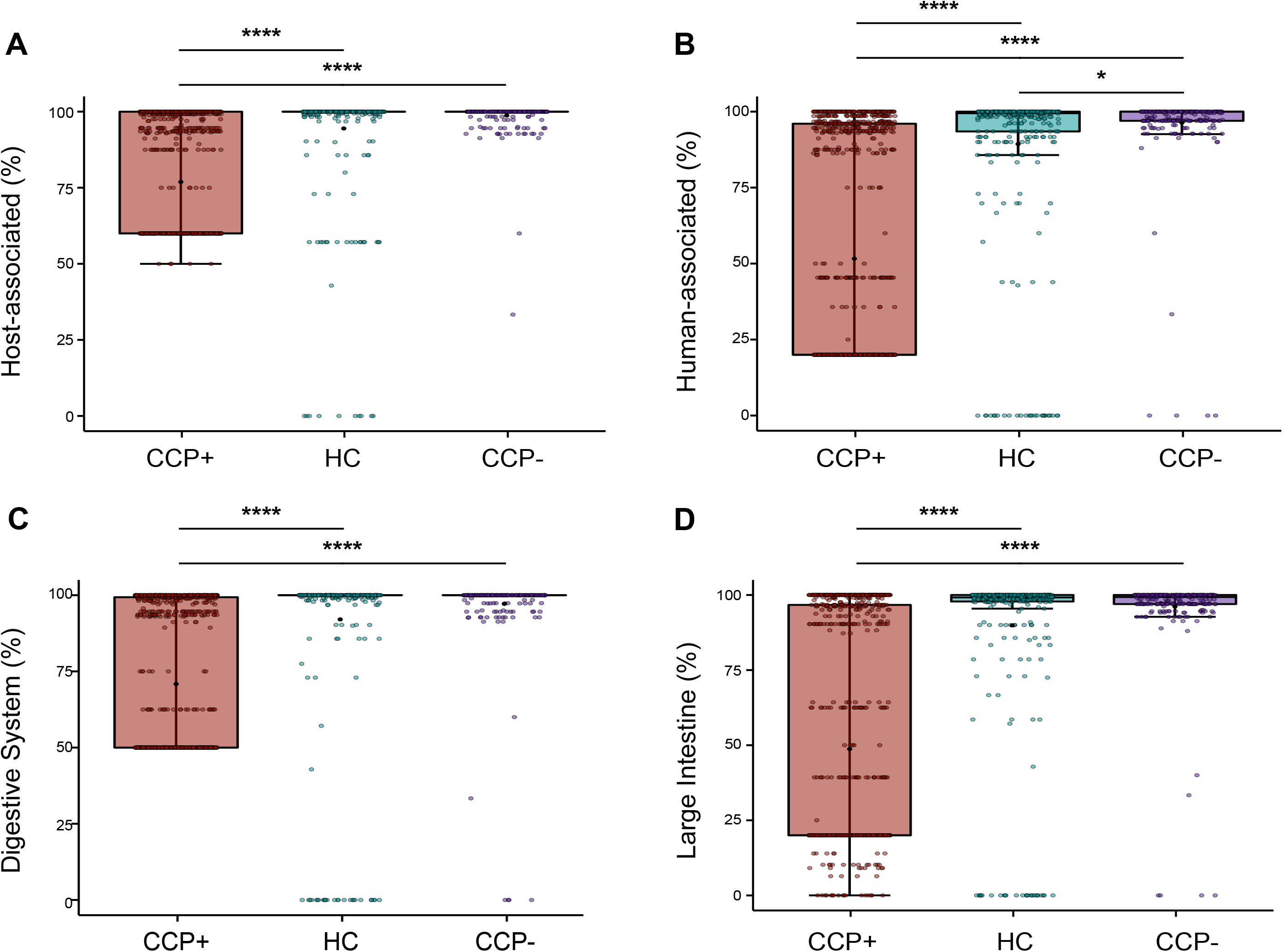
CRISPR spacer host metadata reveal CCP+ phages represent greater variability in microbial host ecology. Phage host isolate ecology metadata was compiled from JGI/GOLD v7.0 and broken down by Ecosystem, Ecosystem Category, Ecosystem Type, and Ecosystem Subtype distributions accordingly for all CRISPR spacers identified within our list of 660 phages. (A) Ecosystem Distribution showing the percent host-associated spacers calculated for each contig based on cohort distribution. (B) Ecosystem Category distribution showing the percent human-associated spacers. (C) Ecosystem Type distribution showing the percent of contigs that contain spacers originating from the digestive system. (D) Ecosystem Subtype showing the percent of contigs that contain spacers originating from the large intestine microenvironment. Cohort distributions based on these metadata revealed a disproportionate distribution of CRISPR spacers among samples originating from CCP+ individuals when compared to CCP- or HC groups. Statistical significance was determined using pairwise Wilcoxon rank sum tests for comparisons between the three groups, using the Benjamini-Hochberg correction for multiple testing comparisons (* *p* = 0.023, **** *p* < 2 × 10^−16^).

These compositions of multiple CRISPR spacer ecosystem distributions reveal homogeneity among phages derived from HC and CCP-samples, and indicates more dysbiotic communities across CCP+ samples, suggesting that CCP+ individuals harbor disparate phage communities that are more likely to originate from non-host associated sources. The putative origins of these phages are related to environmental metadata of CRISPR spacers in the JGI GOLD database describing the origin of bacterial DNA samples across ecologically diverse biomes worldwide [52]; and increased heterogeneity in the CCP+ phages suggests a condition-dependent host intestinal environment that maintains diversity. At the highest Ecosystem classification level, with only three unique classification terms, these non-host associated sources that are more prevalent in the CCP+ group, correspond to a higher degree of spacers matching organisms originating from environmental and/or engineered habitats as archived in GOLD (Figure S5). The ecosystem distributions of Category, Type, and Subtype have 43, 126, and 146 unique terms for each classification level respectively, indicating multiple possible combinations for organism habitats. Thus, our analysis of GOLD metadata for all phages with predicted host isolates within our study reveals divergent habitat origins for CCP+ derived contigs.

### Quantitative read mapping reveals differentially abundant contigs despite sample cohesiveness

We next asked whether certain phage community members are present in different abundances among the members of the cohort at-risk for rheumatoid arthritis compared to healthy controls. To assess differences between phages among the sample groups, we used a viral read recruitment strategy whereby VLP reads from all samples were mapped to the 660 curated contigs [23, 48]. Using read count matrices for all contigs as input in the DEseq2 statistical package for differential analysis of comparative count data [53], we analyzed three pairwise comparisons for over- or under-abundant viral contigs (Figure 5). Initial comparisons of the normalized and log-transformed count matrices were performed to evaluate the experiment-wide trends across all samples. Principal component analyses reveal minimal variance explained by the first two principal components for CCP+ vs HC samples (Figure 5A), CCP-vs HC samples (Figure 5B), and CCP+ vs CCP-samples (Figure 5C), indicating that total sample community signatures cannot be readily differentiated based on at-risk or healthy control cohorts. We further explored the sample similarities by comparing Euclidian sample-to-sample distances of the regularized log-transformed count matrices. Hierarchical clustering of sample-to-sample distances did not reveal any discernable clustering for CCP+ vs HC samples (Figure 5D), and only minimal similarities between two CCP-samples when compared to the HC (Figure 5E) and CCP+ (Figure 5F) groups, suggesting general sample cohesiveness between cohorts.

**Figure 5.**
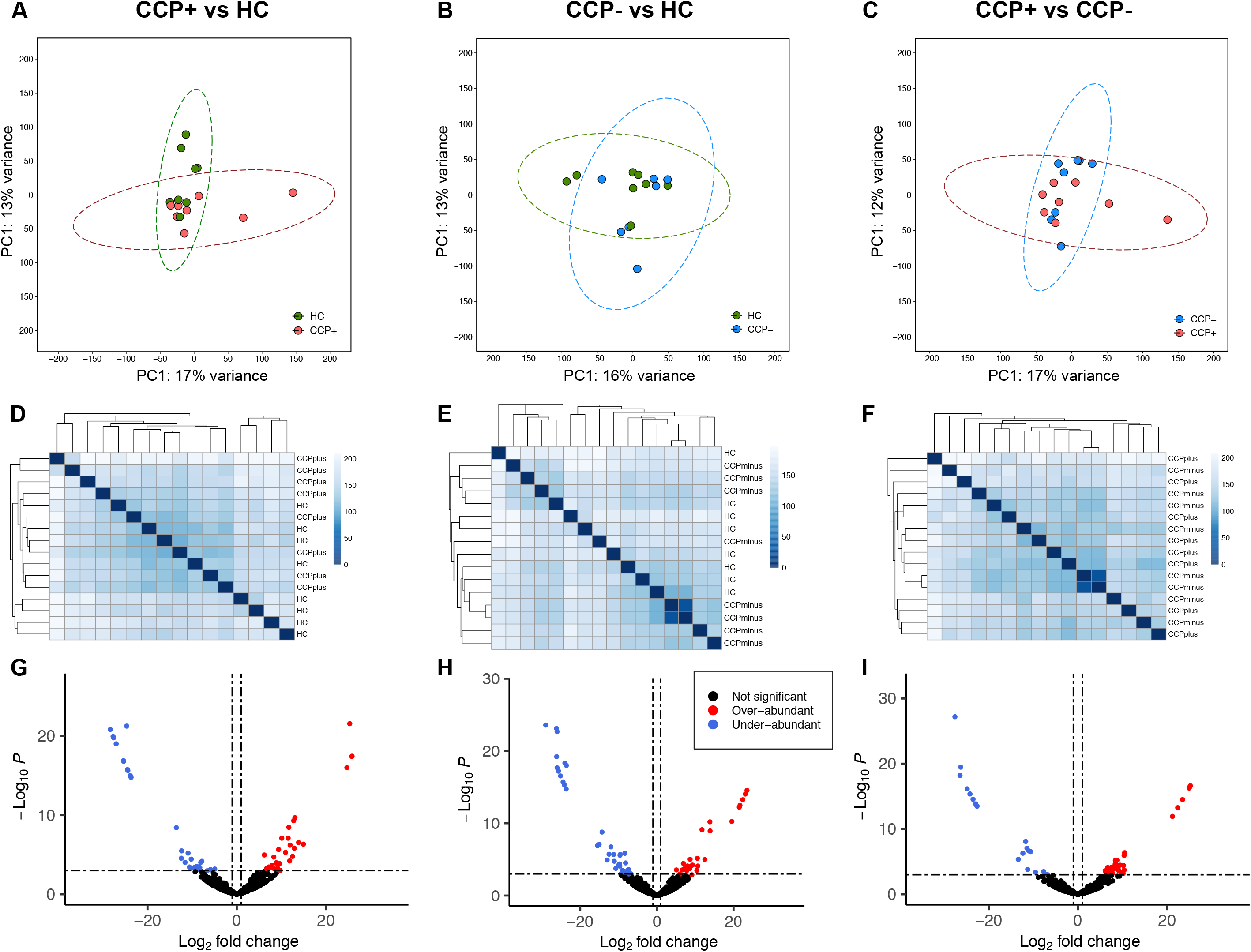
Quantitative read mapping exposes differentially abundant contigs despite sample cohesiveness. Quantitative read mapping of all VLP read sets to the final curated 660 phages reveals contig-contig dissimilarities despite minimal sample-sample variance or intra-sample hierarchical clustering. Differential abundance calculations were carried out within the DESeq2 package by way of 3 pairwise comparisons: CCP+ vs. HC, CCP-vs. HC, and CCP+ vs. CCP-. (A, B, C) Analyses of the first and second principal components for sample-to-sample exploratory analyses revealed minimal variance explained across all comparisons. (D, E, F) Euclidian distances for sample-sample read-based coverages were used for hierarchical clustering across all pairwise comparisons reveal minimal clustering based on sample type. (G, H, I) Volcano plots reveal 9%, 10%, and 8% of contigs included in our analysis are differentially abundant respective to CCP+ vs. HC, CCP-vs. HC, and CCP+ vs. CCP-group-based comparisons of specific contig community members.

We next analyzed specific members of the intestinal phage community, considering the rationale that samples with complex communities are better explored at the level of each unique member [33]. Visualization of the principal components incorporating the viral identification metrics used in the VIBRANT neural network for our 660 curated contigs shows minimal differentiation among phage scaffolds based on scaffold quality (Figure S6A) or predicted phage state (i.e., lytic or lysogenic) (Figure S6B), although fragmentation of smaller sized contigs is evident for both analyses. Further, grouping of contigs at the sample type level does not differentiate any specific cluster (Figure S6C), which is consistent with the minimal variance observed at the sample level (Figures 5A, 5B, and 5C). Finally, we assessed the differential abundance of read recruitment counts for the set of 660 contigs and estimated fold changes based on the negative binomial generalized linear model provided by DESeq2 [53]. Using thresholds of log2-fold change greater than 1 or less than -1 (equivalent to fold change of ± 2) and Benjamini-Hochberg adjusted *p*-values < 0.001, we identified a total of 178 differentially abundant contigs (27% of the 660 phages) across three pair-wise abundance comparisons. For CCP+ vs HC samples a total of 59 contigs (30 over- and 29 under-abundant) (Figure 5G), for CCP-vs HC a total of 66 contigs (27 over- and 39 under-abundant) (Figure 5H), and for CCP+ vs CCP-a total of 53 contigs (27 over- and 21 under-abundant) (Figure 5I) passed our thresholds for significance. This suggests that there are unique changes in select phage abundances from the intestinal viromes of individuals at risk for RA, and that these changes are more nuanced than sample-based community associations can reveal. These data indicate that these cohort groups represent minimal sample-sample variation, but may provide clues related to detection of biomarkers via specific community members. The top phage contigs associated with either CCP+ or CCP-individuals were *Clostridium scindens* (Lachnospiraceae) and *Actinomyces oris* (Actinomycetaceae), respectively, over-abundant at log_2_ fold changes of 25.9 and 23.5 compared to the healthy control samples.

A comparison of the bacterial relative abundances via 16S amplicon sequencing confirmed an expansion of Lachnospiraceae bacteria among samples originating from CCP+ individuals (Figure S7A). This confirms, in part, observations of over-abundant Lachnospiraceae-targeting phage contigs for the CCP+ but not CCP-cohorts (Figures 6B and 6C). The bacterial composition across all cohorts was relatively even in terms of richness (Figures S7B and S7C), evenness (Figure S7D), and species diversity (Figure S7E). Conversely, phage host abundances in the CCP-cohort relative to healthy controls were not correlated to a family-level differentiation in bacterial taxa relative abundance.

**Figure 6.**
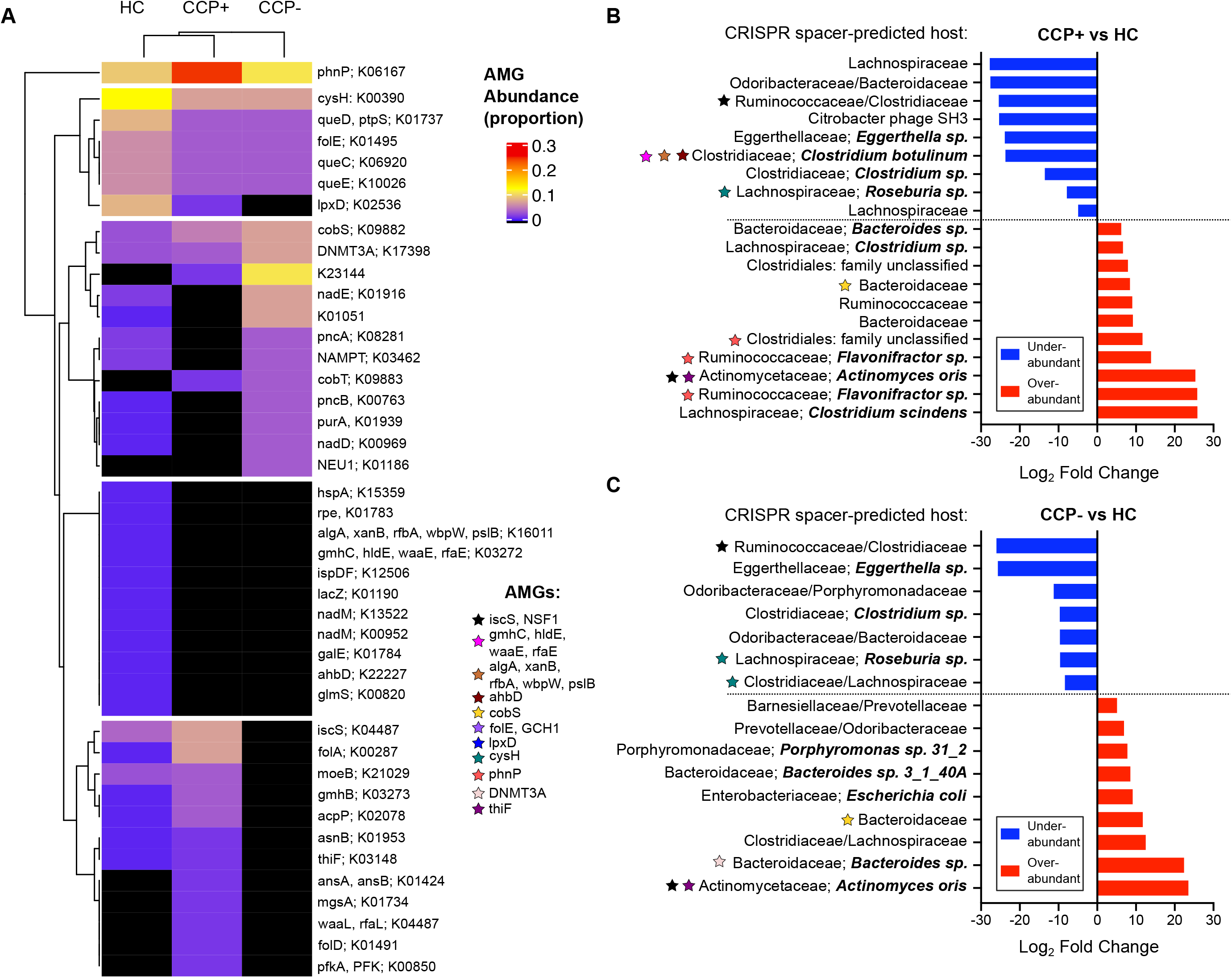
Phage auxiliary metabolic gene abundances highlight cohort-associated disparities in potential metabolic function. AMGs were identified within the VIBRANT algorithm, based on screening 2,835 auxiliary metabolic genes with KEGG Orthology annotations (March 2019 KEGG release, ftp://ftp.genome.jp/pub/db/kofam/archives/2019-03-20/). (A) Total counts per KEGG Pathway were used normalize relative abundance of AMGs per sample, which were clustered using the ComplexHeatmap package in R. Areas in black indicate no AMG hits were present for the entire cohort for the 660 contig samples. (B) Differentially abundant contig for the CCP+ to HC pairwise comparison, visualizing only the contigs which had CRISPR spacer-predicted hosts. Color-coded stars belong to a list of AMGs and indicate association with the contig they are adjacent to. (C) Differentially abundant contigs for the CCP-vs HC comparison.

### Phage auxiliary metabolic gene abundances highlight cohort-associated disparities in metabolic potential

To determine the functional potential and metabolic capabilities within intestinal phages, we quantified AMGs assigned to specific metabolic pathways in the Kyoto Encyclopedia of Genes and Genomes (KEGG) database across at-risk and healthy cohorts. Since their identification as viral drivers of host metabolism [54], phage-encoded AMGs have been recognized as consequential actors that redirect host functional capacities thereby directly influencing local ecology [55, 56]. Analyses of AMGs using VIBRANT and KEGG pathway annotations can provide valuable insights into potentially altered metabolic functions or informative biosignatures for cohort-associated microbial communities [37, 57]. To this end, we assessed our set of curated phage contigs against 2,835 AMGs with KEGG annotations identified as “metabolic pathways” or “sulfur relay system” [37]. Among our 660 phage contigs, 161 (24%) were found to encode at least 1 AMG, with 252 AMGs in total across all samples (Supplemental Table 4). Phages originating from the HC cohort accounted for 131 metabolic signatures, while CCP+ and CCP-had less total AMGs with 77 and 44, respectively (Figure S8A). Among the most represented metabolic categories across all phages, amino acid metabolism and the metabolism of cofactors and vitamins contained 121 and 88 AMGs, respectively, with energy metabolism being the next largest category with 22 AMG hits (Figure S8B). These general pathway results indicate that phages in the intestine presumably affect host metabolism through the consumption of metabolic resources needed for their own biogenesis, as described in phage-host infection studies of model pathogens [58-60] and marine virocells [61].

To further probe all metabolic phage-encoded functions corresponding to our sample cohorts, we assessed all AMG hits for total KEGG pathway abundances. Hierarchical clustering grouped AMGs into 5 distinct metabolic clusters relative to HC and at-risk CCP cohorts (Figure 6A). Among these groups, the gene coding for *phnP* (K06167) stands apart from the others, both in terms of clustering and also for relative pathway abundance (Figure 6A). Among group-associated differences in AMG pathway abundances, there are notable absences among both CCP+ and CCP-individuals. Namely, several clustered transferases such as the mannose-phosphate transferases (*algA, xanB, rfbA, wbpW, pslB*), manno-heptose transferases (*gmhC, hldE, waaE, rfaE*), and the *galE* epimerase and *glmS* transaminase (Figure 6A). Considering the impact of such transferases on bacterial cell wall polysaccharides and biofilm formation [62, 63], these results point to a baseline of phage-driven bacterial surface modifications from HC-derived phages. Conversely, AMGs involved in lipopolysaccharide (LPS) biosynthesis such as the *waaL* O-antigen ligase and the *gmhB* phosphatase are only present in CCP+ phages or at greater abundance in CCP+ phages, respectively, indicating a possible role in immune evasion. Within the CCP-cohort, one of the most abundant AMGs, KEGG orthology entry K23144 encoding for a polyketide sugar transferase important in peptidoglycan biosynthesis is completely absent from the HC cohort and present at lower levels for CCP+ samples. Thus, phage-encoded bacterial surface modifying enzymes such as the sugar transferases and LPS/peptidoglycan biosynthetic genes, are differentially represented across the cohorts in this study, which has implications for bacterial fitness in the intestinal ecosystems and their interactions with the immune system.

We next incorporated the AMG characterization of genomes within our curated set of phages to those that were significantly over- or under-abundant in previous differential abundance analyses (Figures 5G, 5H, and 5I). Among the 20 differentially abundant contigs from the CCP+ vs HC pairwise comparison that contained CRISPR spacer-predicted hosts, 8 of these encoded at least one AMG (Figure 6B). The 9 under-abundant phages in this comparison encode 5 AMGs, including manno-heptose transferases (*gmhC, hldE, waaE, rfaE*), mannose-1-phostphate transferases (*algA, xanB, rfbA, wbpW, pslB*) and *ahbD* AdoMet-dependent heme synthase all together on 1 contig, and *cysH* and *iscS* genes on 2 other contigs (Figure 6B). Among the 11 significantly over-abundant contigs, 3 of these encode the *phnP* phosphodiesterase; 3 phages predicted to infect *Flavonifractor* sp. (Ruminococcaceae) and one predicted to infect Clostridiales bacteria. The remaining AMG found in CCP+-associated over-abundant phages encodes for a cobalamin biosynthesis protein *cobS*, found in marine cyanophages [64], viruses of marine archaea [65], and is considered a core component of marine phage genomes [66], but also ubiquitous in phage genomes that infect *E. coli* [67]. Our identification of a CCP+ over-abundant phage contig targeting *Bacteroides fragilis* and carrying the *cobS* AMG (Figure 6B) reinforces the universal nature of this central AMG that is conserved across hosts and environments [37].

We also identified 16 unique phage contigs with definitive CRISPR spacer-predicted hosts that were differentially abundant and associated with the CCP-cohort (Figure 6C). Within these contigs, 9 are significantly under-abundant compared to healthy controls, with 3 of these encoding AMGs. CCP-associated phages were identified as carrying *cobS, DNMT3A, thiF*, and *iscS* metabolic genes (Figure 6C). Thus, in contrast to CCP+ associated contigs which harbored *phnP* and *cobS* on a combination of Lachnospiraceae, Rumminococcaceae, and Bacteroidaceae targeting phages, CCP-associated phages were identified to target primarily Bacteroidaceae and *Actinomyces oris* and harbor a combination of AMGs.

## DISCUSSION

RA is a complex disease with an unknown etiology that puts a burden on quality of life resulting in a strong societal impact [68, 69]. In addition to multiple epidemiological factors being associated with RA development, including genetic and familial risk, environmental risk factors and biological sex [3], the microbiota remains an important and understudied factor that likely influences RA autoimmunity [70]. Given the widespread occurrence and diversity of phages in the human intestinal microbiota and their impact on intestinal microbial ecology during health and disease [19, 20, 71], we analyzed this previously neglected component of the microbiota as it relates to RA etiopathogenesis. We used shotgun metagenomics to identify intestinal phages of individuals at risk for developing RA and discovered an association of distinct phage communities with RA-specific serology in the at-risk population.

Using three separate database-independent approaches, we describe a collection of 660 phage genomic sequences, their potential metabolic capability, and their differential abundance. Through a combination of CRISPR spacer matching and Markov clustering with other viral metagenomic sequences from diverse environments, we predicted host assignments for 285 or 43.2% of these phages, which is a high level of taxonomic assignments relative to recent reports of approximately 10 – 30% host assignment identification [23, 48, 72]. By analyzing a core set of *de novo* assembled phage contigs paired with taxonomy, we identified differential phage communities associated with the at-risk RA individuals compared to healthy controls, all while adding novel phage-host assignments to previously unidentified intestinal phages [73, 74].

Phage-host assignments were dominated by Lachnospiraceae-targeting phages, some of which were over-abundant in CCP+ individuals. This expansion of phages also correlated with increased abundances of Lachnospiraceae bacteria in the CCP+ cohort compared to either CCP- or the healthy cohort, suggesting a link to this family of Firmicutes and CCP autoantibody production in the human intestine. Interestingly, increased abundance of Lachnospiraceae has been observed in at least two previous studies of intestinal microbiotas in mice during the course of collagen-induced arthritis (CIA) [75, 76]. Considering the precedence for overlap of identified phage contigs from mouse intestines to human-associated intestinal phages [23], the previously-reported increase in abundance of Lachnospiraceae bacteria during experimental arthritis in mice is supported by our findings of increased Lachnospiraceae phage-host interactions in CCP seropositive individuals at-risk for developing RA. To this end, given that the FDR individuals included in this study do not show clinical signs of established RA, our identification of a preclinical cohort with increased Lachnospiraceae phage-host interactions could serve as a biological indicator of disease. Similarly, an expansion of Bacteroidaceae-targeting phages associated with the CCP-cohort was described, which corresponds to a previously observed expansion of Bacteroidaceae bacteria following CIA in mice [75]. In addition to these phages serving as potential biomarkers of disease in humans at risk for RA, our data indicate that Bacteroidaceae and Lachnospiraceae-targeting phages designate a distinction between CCP serology status that may serve as an additional indicator of disease progression and/or future disease severity [77]. Notably, bacteria in the Lachnospiraceae and Ruminococcaceae families have been linked to the pre-diabetic intestinal microbiota and diabetic pathogenesis, while Bacteroidaceae are associated with disease protection in a murine model of diabetes [78]. The identification of cohort-specific phage-host interactions sheds light on potential preclinical biomarkers connecting specific dysbiotic intestinal microbial communities to possible regulation of microbiota-mediated mucosal inflammation [1, 79].

We calculated the differential abundance of curated phages on a contig-to-contig basis to estimate dispersion and fold changes of quantitative read mapping matrices. In doing so, we identified 178 differentially abundant contigs (27% of the total curated list) across three pair-wise cohort comparisons. Among the CCP+ vs HC comparison, we observed over-abundant phages targeting *Clostridium scindens, Flavonifractor* sp., *Actinomyces oris*, as well as other family-level taxonomic assignments. A member of the Lachnospiraceae, *C. scindens* is an intestinal commensal bacterium involved in maintaining homeostatic large intestinal bile acid composition and providing host protection from opportunistic *Clostridioides difficile* blooms [80, 81]. A differential abundance of phage targeting *C. scindens* in the CCP+ at-risk cohort, may have implications for bile acid dysmetabolism in these individuals, which has consequences for inflammatory bowel diseases [82, 83]. Differential abundance of phages in the CCP-cohort revealed several phages targeting Bacteroidaceae and *Bacteroides* species, bacteria involved in multiple reactions of bile acid metabolism promoting host metabolic health [84, 85]. Recent phage-*Bacteroides* interactions have described the influence of phage BV01 in reducing *Bacteroides* bile acid metabolism [86], which has implications for the impact of phages on mammalian gut metabolic function. Our findings suggest individuals at risk for RA harbor divergent communities of phages with potential to alter intestinal metabolic potential through either reduction of key bacterial species and thus reducing endogenous metabolic function, or through the phage-derived introduction of specific AMGs.

Changes to the intestinal metabolome can lead to compositional microbiota transitions that in turn impact host nutrient uptake and immune homeostasis [87]. Considering that manipulations of microbial metabolic pathways in the intestine can influence inflammation and dysbiosis [88], our identification of phage communities with differential abundances of encoded AMGs points to divergent metabolic landscapes associated with at-risk RA cohorts. A majority of the AMGs identified in our analysis make up a group of 14 genes conserved across many environments [37], indicating their functional importance in core metabolism. We were surprised to identify three phages that were over-abundant in the CCP+ cohort (3 of 11 in total), three with *Flavonifractor sp*. predicted hosts and one Clostridiales-targeting phage, encoding the *phnP* phosphodiesterase. Encoding a phosphoribosyl 1,2-cyclic phosphate phosphodiesterase, *phnP* accounts for 10% of the total AMGs represented in our phage genomes, and is differentially abundant among the CCP+ cohort samples. While *phnP* is one of 14 genes considered to be globally conserved across multiple environments [37], it is the only gene among AMGs in our analysis that is the lone representative of its pathway. The PhnP phosphodiesterase, part of a 14-gene operon originally described in *Escherichia coli*, plays a crucial intermediary role in the carbon-phosphorous lyase pathway by degrading a dead-end cyclic phosphonate byproduct [89]. The uniform presence of *phnP* across phages derived from at-risk and healthy cohorts (Figure 6A), suggests phage-driven organophosphonate degradation, which is fundamental for bacteria in diverse environments [90].

Phosphonate degradation is important for phosphorus assimilation in enteric bacteria [91], although phosphonate metabolism has not been described for *Flavonifractor* species and a *phnP* homolog is not available for this genus in the KEGG database (K06167). In a recent study characterizing microbiota KEGG orthologs as predictors of methotrexate responsiveness for RA treatment, a gene in the phosphonate transport system, *phnC* (K02041), exhibited high median random forest importance as a predictor of drug response in new-onset RA subjects [92]. The contribution of the phosponate metabolic pathway in bacteria and phages, will require further exploration in the context of RA pathogenesis and treatment. However, it is possible that these phage-encoded metabolic products are supplementing phosphorous uptake among Ruminococcaceae and Lachnospiraceae bacteria that predominate in CCP+ individuals prior to RA clinical symptoms. Our analysis is limited in that we did not measure a longitudinal progression of microbial metabolic pathways in these human samples, yet these metabolic associations warrant further investigations into causality and the potentially cascading effects on interbacterial interactions [93].

Our results point to divergent communities of phages with multiple bacterial host targets that group according to anti-CCP serology in individuals predisposed to developing RA. These at-risk individuals who develop seropositive RA, a disease manifestation that is more severe [94] and less responsive to treatment [95], endure a prolonged asymptomatic period before pathological early RA develops in those who are at a higher disease susceptibility in the preclinical RA state [1]. Current approaches for RA diagnosis rely in large part on anti-CCP serology which has up to 93% specificity but as low as 67% sensitivity (for the CCP3.1 assay used here) [39], indicating that a negative result does not preclude current or development of clinically apparent RA. Phage community composition analyses may complement existing diagnoses for RA, considering that intestinal phages can play important roles in immune tolerance, mucosal immunity, and microbial homeostasis [96]. Given that phage community alterations have been shown to precede autoimmunity development in children at risk for developing type 1 diabetes [26], phage community structure should be considered as a biomarker for diseases such as RA that are influenced by non-genetic microbial factors [19]. To that end, we have characterized the intestinal viromes of RA at-risk individuals corresponding to anti-CCP serology status. Furthermore, we calculated species-specific phage-host interactions and identified over-abundances of *C. scindens* and *A. oris* targeting phages in CCP+ and CCP-individuals, respectively. Divergent metabolic profiles evident by differential abundance of AMG-encoding phages in both conditions warrant further interrogation during models of RA-like disease. Future work should investigate the potential of phages in a murine CIA model to determine the influence of RA-associated phages with immunomodulation and inflammatory disease progression. Our multifaced approaches for phage prediction and phage host assignments hold promise to better ascertain the occurrence and diversity of the virome and the identification of key phages influencing the microbiota and individuals at risk for developing RA autoimmune disease. This RA-focused study implicating specific phage populations could open new avenues to assess the basis for phage implication in other microbiota dysbiosis-associated diseases.

## RESOURCE AVAILABILITY

### Lead Contact

Further information and requests for resources and reagents should be directed to and will be fulfilled by the Lead Contact, Breck A. Duerkop.

### Materials Availability

This study did not generate new unique reagents.

### Data and Code Availability

The VLP and whole metagenome DNA sequencing reads as well as the final curated phage contigs generated in this study are available at the European Nucleotide Archive under the Study titled “Intestinal VLP reads and predicted phage contigs for at-risk RA individuals” (accession numbers PRJEB42612 and ERP126498). The VLP and whole metagenome raw unmapped read sets are available for each of the 25 individual samples included in this study and are available under the Study Primary Accession PRJEB42612. The 660 curated contigs are compiled in a multifasta file deposited as Sample SAMEA7856466 under the same Study PRJEB42612.

## EXPERIMENTAL MODEL AND SUBJECT DETAILS

### Study Subjects and Fecal Samples

Fecal samples were obtained from individuals recruited for the SERA (Studies of the Etiology of Rheumatoid Arthritis) initiative, aimed at understanding the mechanisms that prelude the preclinical development of RA. SERA is a multicenter prospective cohort study that has identified first-degree relative (FDR) probands defined as a parent, full sibling, or offspring of individuals with diagnosed clinical RA [38]. FDR probands were evaluated in extensive clinical research visits, longitudinal follow-ups, and autoantibody testing to determine CCP status [38]. FDR probands were split into cohorts dependent on serum CCP levels, with 100% of subjects in the CCP+ cohort positive and 0% of subjects in either CCP- or HC (Healthy Control) cohorts testing positive. Healthy control subjects were recruited and included in this study as described previously [97]. The present study consisted of 25 subjects split into 3 cohorts, of which 8 were CCP+, 8 were CCP-, and 9 were HC. Ethical approval for this study was obtained from the University of Colorado Multiple Institutional Review Board (COMIRB) study numbers 01-675 (primary) and also 13-2606 and 14-1751. COMIRB Protocol 01-675 included informed consent with HIPAA authorization for stool sample collections. Stool samples were obtained independently by SERA study participants and returned within 1 week of their original visit. Samples were stored in aliquots at -20°C until processing.

## METHODS

### Extraction of Fecal Whole Metagenome and VLP DNA, Library Preparation and Sequencing

Whole metagenome and VLP fraction DNA were isolated as described previously [98], with some modifications as follows. For all samples, 0.1 g of human stool was homogenized in 8 mL salt magnesium plus (SM+) buffer [99] and 0.5 ml of homogenate was transferred to a BashingBead Lysis tube (Zymo) and designated as the whole metagenome sample. Whole metagenome DNA was extracted using a ZymoBIOMICS DNA kit (Zymo) following the manufacturer recommended protocol. VLPs were clarified from the remaining 7.5 mL of sample by three successive centrifugation steps (3200g for 10 min, 3200g for 10 min, 7800g for 10 min), and the supernatant was filtered through a 0.45-µm PVDF filter membrane. VLPs were precipitated by adding 0.5M NaCl and 10% wt/vol PEG8000 and incubating on ice at 4ºC overnight, followed by centrifugation (7800g for 20 min). VLP pellets were resuspended in 400 µL SM+ buffer and treated with 40 µL DNase buffer (10 mM CaCl_2_, 50 mM MgCl_2_), 25 units DNase, and 15 units RNase for 1 hr at 37ºC. VLPs were further treated with 50 mg/mL proteinase K and 0.5% SDS for 30 min at 56ºC before addition of 100 µL phage lysis buffer (4.5 M guanidiniumisothiocyanate, 44 mM sodium citrate pH 7.0, 0.88% sarkosyl, 0.72% 2-mercaptoethanol) and incubated for 10 min at 65ºC. VLP DNA was precipitated and extracted with an equal volume of phenol/chloroform/isoamyl alcohol 25:24:1, spun at 7800g for 5 min, and further extracted with an equal volume of chloroform. VLP DNA was precipitated with 0.3M NaOAc (pH 5.2) and an equal volume of isopropanol, washed with ice-cold 70% ethanol, and resuspended in sterile water.

### Metagenomic DNA Sequencing

VLP and whole metagenomic DNA was sequenced using the Illumina NovaSEQ 6000 platform with paired-end 150-cycle sequencing chemistry. DNA libraries were amplified using the Ovation Ultralow System v2 (Nugen, part no. 0334) library preparation kit including 12 cycles of amplification. TruSeq adapters (Illumina) were used for multiplexing. Libraries were quantified using a Qubit and quality was measured using a Tapestation. All library preparation, quantification, quality assessment and control, were performed by the University of Colorado Anschutz Medical Campus Genomics and Microarray Core.

### 16S rRNA Amplicon Sequencing and Analysis

16S rRNA gene analysis was performed using fecal samples that were processed for isolation of whole metagenomic DNA using a ZymoBIOMICS DNA kit (Zymo) and stored at -80°C. Amplicons of the 16S rRNA gene V4 region were amplified using Earth Microbiome Project primers 515F and 806R [100] with custom barcodes. Samples were sequenced on the Illumina MiSeq platform with paired end 250 bp reads using bTEFAP technology [101] by MR DNA (Molecular Research LP, Shallowater, TX), and processed using mothur v.1.44.0 [102]. Sequenced reads, which averaged 607,915 ± 112,641.7 per sample, were demultiplexed, assembled as contigs, and processed to remove chimeras and erroneous sequences per the Kozich protocol [103] and mothur MiSeq SOP (https://mothur.org/wiki/miseq_sop/ accessed 07/16/2020). Sequences were aligned to the Greengenes core reference alignment for taxonomy using the 2013 release (gg_13_8_99) [104]. Sequences were differentiated into amplicon sequence variants (ASVs) using the make.shared command, resulting in a total of 8,108,071 sequences. Subsampling was performed using 186,745 sequences, which corresponded to the smallest sample in our dataset. Diversity measurements were performed using mothur calculators to estimate community richness (Chao1 estimator), community evenness (Shannon evenness), and community diversity (inverse Simpson index).

### Decontamination and Read Processing

Metagenomic reads were decontaminated and trimmed as previously described [23] using BBMap short read aligner v38.56 [105]. Briefly, raw reads were mapped to the internal Illumina phage genome control phiX174 (J02482.1), human reference genome (hg38), and potential laboratory contaminants including mouse genome (mm10), *Enterococcus faecalis* V583 genome (NC_004668.1), *E. faecalis* OG1RF genome (NC_017316.1), and *E. faecalis* phage VPE25 (LT546030.1) using the bbsplit algorithm with default settings. Unmapped reads were trimmed of adapter sequences, with low quality reads and reads of insufficient length removed using the bbduk algorithm with the following parameters: ktrim = lr, k = 20, mink = 4, minlength = 20, qtrim = f, as previously described [23].

### Metagenomic Assembly

Decontaminated and trimmed R1 and R2 reads were interleaved using the fq2fa --merge command from the IDBA-UD package [106]. Whole metagenome and VLP assemblies were performed using the MEGAHIT assembler v1.2.7 [107] using the default setting plus the following flags: --presets meta-large (--k-min 27 --k-max 127 --k-step 10) for large and complex metagenomes.

### Quantitative Read Mapping and Construction of the Curated VLP Contig Database

VLP reads were assembled into 25 individual sample sets, corresponding to the 25 individual fecal samples included in our study. All contigs resulting from MEGAHIT assembly were filtered to a minimum length of 5kb, resulting in a pool of 80,762 total contigs from all samples. Three separate independently published methods were employed to identify putative phages from the pooled set of contigs over 5kb in length. First, the P/M read mapping approach was used whereby each sample’s VLP and whole metagenome reads were mapped to their corresponding assembled contigs, using BBMap as previously described [23]. After pooling, the top 100 largest ratios of VLP reads to whole metagenome reads for all 25 read-mapping sets for each sample were identified and pooled. Redundancy was removed using cd-hit-est at an identity threshold of 95% resulting in 2117 unique contig sequences. Next, as a separate method, putative phages from the 80,762 contigs were identified by searching for viral protein family (VPF) hits, as previously described [41]. Separate filters were applied for VPF hits calculated in relation to total genes, microbial genes, and percent non-viral Pfams. 2,902 contigs were identified that contained 5 or more VPF hits and with non-viral Pfam hits below 20%. 263 contigs were identified with 5 or more VPF hits, with more viral gene content than microbial genes per contig, and 644 contigs were identified with 2 – 4 VPF hits and 0 microbial gene hits. Finally, 976 contigs were identified with only 1 VPF hit per contig, and were included regardless of microbial gene content. The third and final independent phage contig identification method used was VIBRANT v1.2.1 [37], a neural network machine learning algorithm that identifies viral protein signatures. VIBRANT identified 4,758 unique phage contigs. After combining these three independent approaches used to identify unique sets of phages, all sets were combined and the overlapping 660 contigs were used for analysis as the curated contig set. To assess contig completion and contamination levels, CheckV v0.6.0 was used with standard operating parameters.

### Differential Abundance Analyses

To calculate differential abundance in pairwise analyses, we first generated read mapping count matrices by mapping all VLP reads to the curated contig set of 660 contigs. The bbmap algorithm from the BBMap suite of tools was used with the following parameters: ambiguous = random, qtrim = lr, minid = 0.97. Total raw read counts were aggregated per contig and assembled into 25 count matrices for all samples, which were then used as input for DESeq2 v1.20.0 [53] running in R version 3.6.3 for comprehensive differential abundance analysis. Raw un-normalized read count coverage values were used to compare fold changes across three pairwise comparisons: CCP+ vs. HC, CCP-vs. HC, and CCP+ vs. CCP-groups. The standard workflow for differential analysis within DESeq2 was used, producing logarithmic fold-change values incorporating Wald tests for *p*-value calculations and the Benjamini-Hochberg multiple testing correction for the adjusted *p*-value. In total, 178 phage contigs from our set of 660 were found to be differentially abundant using thresholds of log_2_ Fold Change < -1 or > 1 and adjusted p-value < 0.001.

### VLP Clustering, Phage Host Matching, and AMG Identification

Clustering of all viral contigs within the RA virome described in this study was performed using two lists of contigs, the total 4,785 viral sequences identified by all filters of the VPF method, as well as the final curated set of 660 contigs. First, all 4,785 sequences were screened against the most recent iteration of the public viral database IMG/VR v3.0 [44] using blastn with 95% sequence similarity over 85% of each 1kb region of the contig, which resulted in 19,892 viral sequences. Then, a total of 24,926 sequences were screened against each other using blastn with the same parameters and omitting duplicate hits. Markov clustering of these 9.4 million connections resulted in a total of 1,193 clusters encompassing 22,306 total sequences. Overall, 2,420 of the 4,785 total RA virome sequences were clustered into 1,184 clusters. Of these clusters, 41 contained a reference viral isolate, 1,037 contained another metagenomic viral contig from IMG/VR, and 106 were identified as originating from RA metagenomic sequencing projects. Lastly, clustering was also calculated for the 660 curated viral sequences, which resulted in 266 individual clusters containing 336 (roughly 48% of curated set) unique sequences. Phage host assignments were determined via bacterial CRISPR spacer matching as previously described [23], requiring at least 93% sequence identity match over the entire spacer length and allowing for up to 2 mismatches. Of our 660 curated contig list, 207 (31.4%) had CRISPR spacers matching reference isolates therefore leading to host predictions for a third of our final contigs. VIBRANT v1.2.1 was used to identify auxiliary metabolic genes (AMGs) according to KEGG metabolic pathway annotations. VIBRANT annotates using VOG, Pfam, and KEGG databases; therefore, if the best HMM hit is to the KEGG database and also if the annotation is in a metabolic pathway, the hit gets called as an AMG.

### Data Visualizations

Various R packages were used, including DESEq2, ggplot2, ComplexHeatmap, pheatmap, corrplot, RColorBrewer, and EnhancedVolcano. Graphpad Prism v8.4.3 was used for all supplemental calculations. Lastly, SankeyMATIC (https://github.com/nowthis/sankeymatic) and meta-chart (https://www.meta-chart.com/venn) were used to create the Sankey and Venn diagrams depicted in Figure 1, respectively.

## SUPPLEMENTAL INFORMATION

**Figure S1. Overview of methods for VLP isolation and phage identification from sequencing reads**. (A) Individual stool samples were homogenized and split into P and M subsamples for generating VLP and whole metagenome DNA, respectively. (B) Total sequencing reads generated per sample for each P and M read sets, after quality control and decontamination. (C) Total assembled contigs with length greater than 5kb generated per sample for each P and M read sets. (D) Overview of the computational pipeline used to identify phages; from short-insert pair end read sets averaging approximately 75M read pairs per sample, to the 80,762 *de novo* assembled contigs greater than 5kb in length, and the three independent methods for phage identification (P/M, VPF, VIBRANT).

**Figure S2. Estimation of contig completeness by CheckV**. Distribution of contig lengths across contig quality categories according to the MIUViG standards. Contigs derived from the (A) P/M ratio method of phage identification, (B) the VPF method, (C) VIBRANT algorithm, and finally (D) the curated contig list. Boxplots depict the following five summary statistics: median, lower and upper hinges corresponding to the first and third quartiles, and two whiskers corresponding to 1.5 times the interquartile range between the first and third quartiles. Points beyond the whiskers correspond to outlier points.

**Figure S3. Lifestyle and morphology distributions of curated phage contigs**. (A) Total contigs per sample among the three groups, divided according to infection mechanism (lytic vs. lysogenic) as determined by the VIBRANT algorithm. (B) Relative abundances of phage lifestyles as determined by the VIBRANT algorithm. In total, for our 660 predicted phages, 467 (70.8%) are classified as lytic and 193 (29.2%) are classified as lysogenic by VIBRANT. (C) Viral taxonomy of all contigs per sample including the top four morphotypes: Siphoviridae, Myoviridae, Podoviridae, and Microviridae. (D) Relative abundance of all viral morphotypes identified for all 660 phages. Viral taxonomy was determined using a custom database described in this preprint by Kieft et al. (2020; bioRxiv preprint doi: https://doi.org/10.1101/2020.08.24.253096).

**Figure S4. Clustering distribution of singletons and viral groups**. (A) Total singletons, viral contigs that did not cluster with any other genome, per sample and RA cohort group. (B) Distribution of total viral clusters in relation to the number of viral genomes clustered within each group.

**Figure S5. CRISPR spacer host metadata distribution of environmental and engineered derived phages per cohort**. Phage host isolate ecology metadata was compiled from JGI/GOLD v7.0 at the highest Ecosystem classification level for all CRISPR spacers identified within our list of 660 phages. Data is presented as percent of spacers per contig whose metadata is designated as originating from (A) environmental or (B) engineered environments, distributed across the three RA cohort groups. Statistical significance was determined using pairwise Wilcoxon rank sum tests for comparisons between the three groups, using the Benjamini-Hochberg correction for multiple testing comparisons (* *p* = 0.011, **** *p* < 2 × 10^−16^).

**Figure S6. Principal component analyses based on quality, predicted phage lifestyle, and sample cohort**. Principal components for the final curated set of 660 contigs derived from the VIBRANT phage identification program categorized by (A) contig quality, (B) phage lifestyle, and (C) cohort group. Total identified open reading frames were incorporated in analyses in (A) and (B), showing a greater dispersion of smaller sized contigs and a consensus grouping of bigger contigs.

**Figure S7. Analysis of bacterial family diversity from fecal samples based on 16S sequencing and analyzed using mothur**. (A) Relative abundances of bacterial families based on ASV binning reveals a significant difference in Lachnospiraceae bacteria originating from CCP+ fecal DNA samples. Unpaired nonparametric Mann-Whitney tests were used to compare ranks, revealing *p* values of 0.0464 comparing CCP+ to HC individuals. Community richness was measured by the standard observed richness calculator in mothur (B) as well as the Chao1 richness estimate (C). Community evenness was measured using the Shannon index (D), and community diversity was measured using the inverse Simpson index (E). No statistically significant differences were observed among any of the above calculators using nonparametric tests of significance among the three groups.

**Figure S8. Distribution of auxiliary metabolic genes found on curated contigs**. (A) A total of 252 AMGs were discovered among our 660 phages, distributed across the three cohorts. (B) AMGs were categorized predominantly as belonging to amino acid and cofactor/vitamin metabolism categories.

## ACKNOWLEDGEMENTS

The authors thank all participants in the SERA studies at the University of Colorado and elsewhere. We thank Rodolphe Barrangou for comments and suggestions to improve this manuscript. We thank Monica Ransom and Katrina Diener from the University of Colorado Anschutz Medical Campus Genomics and Microarray Core for performing Illumina library preparation and sequencing. The following funding sources supported this work: NIH U01AI101981 (V.M.H., K.A.K.), NIH U01AI130830 (B.A.D.), startup funds from the University of Colorado School of Medicine (B.A.D.), NIH Research Training in Rheumatology T32AR007534 (M.R.M.), Rheumatology Research Foundation Future Scientist Award (M.E.C.).

## AUTHOR CONTRIBUTIONS

Conceptualization, M.R.M, D.P., A.C., K.A.K., and B.A.D.; Methodology, M.R.M., D.P., K.K., A.C., J.A.S., M.L.F., M.K.D., and B.A.D.; Investigation, M.R.M, D.P., K.K., A.C., M.E.C.; Sample Procurement, M.E.C., J.A.S., M.L.F., and M.K.D.; Visualization, M.R.M, D.P. and K.K.; Writing – Original Draft, M.R.M and B.A.D.; Writing – Review & Editing, M.R.M, D.P., K.K., A.C., M.E.C, A.S., K.D.D., V.M.H., K.A.K. and B.A.D.; Funding Acquisition, V.M.H. and B.A.D.; Resources, B.A.D.; Supervision, B.A.D., V.M.H., K.D.D., K.A., A.S. and K.A.K.

