## Supplementary figures and images for "Individuals at risk for developing rheumatoid arthritis harbor differential intestinal bacteriophage communities with distinct metabolic potential"

### Supplemental Figure 1

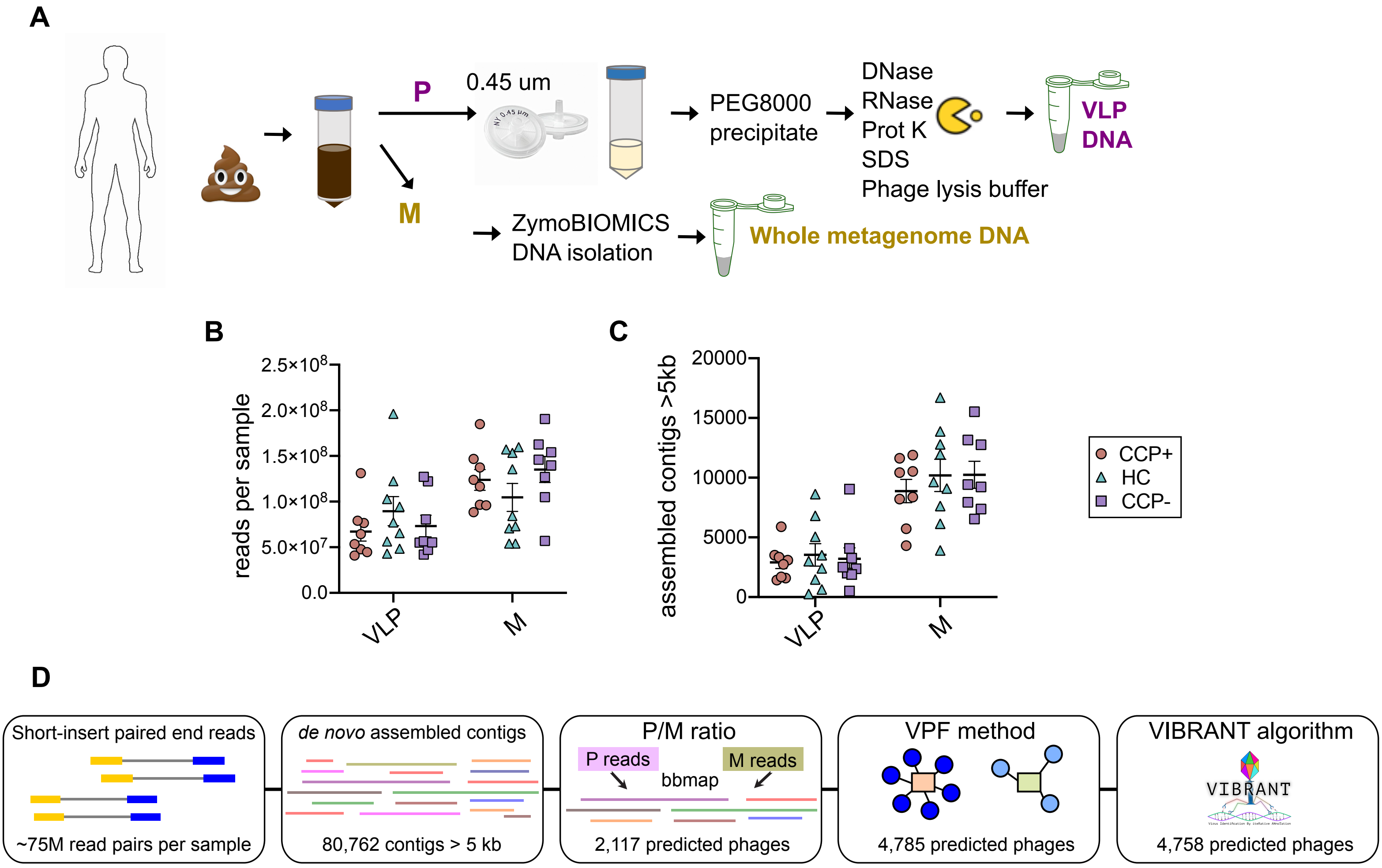

### Supplemental Figure 2

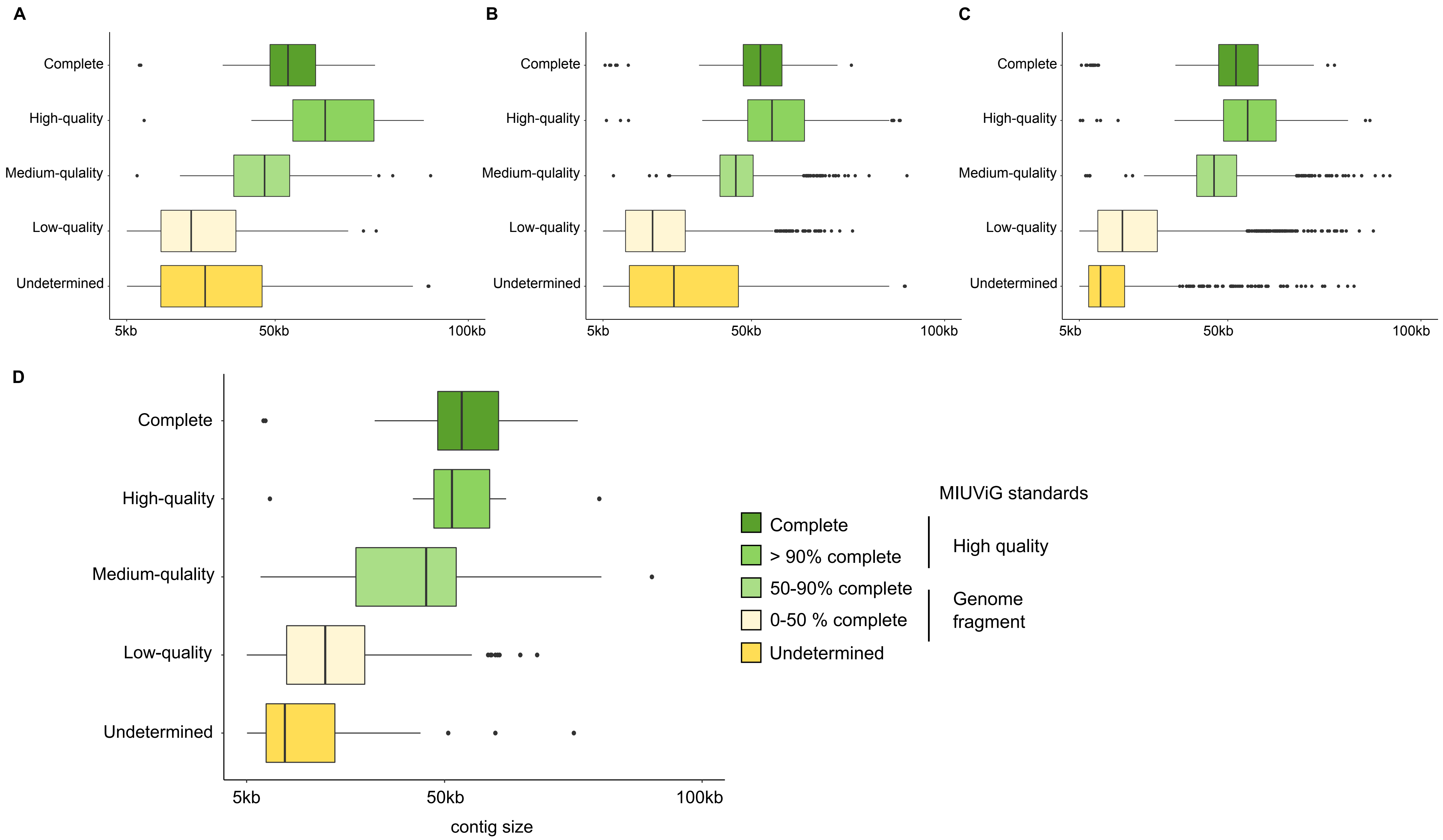

### Supplemental Figure 3

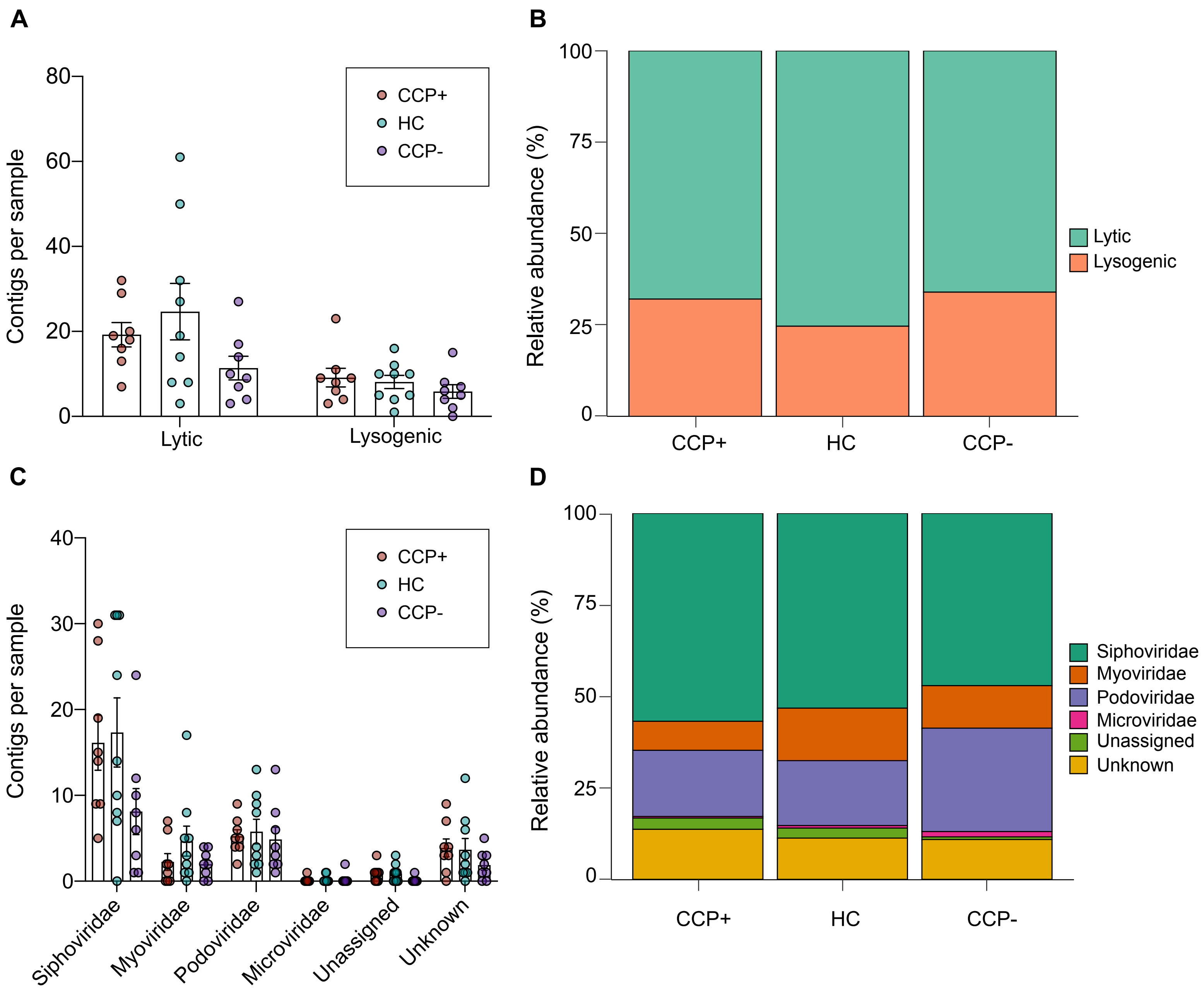

### Supplemental Figure 4

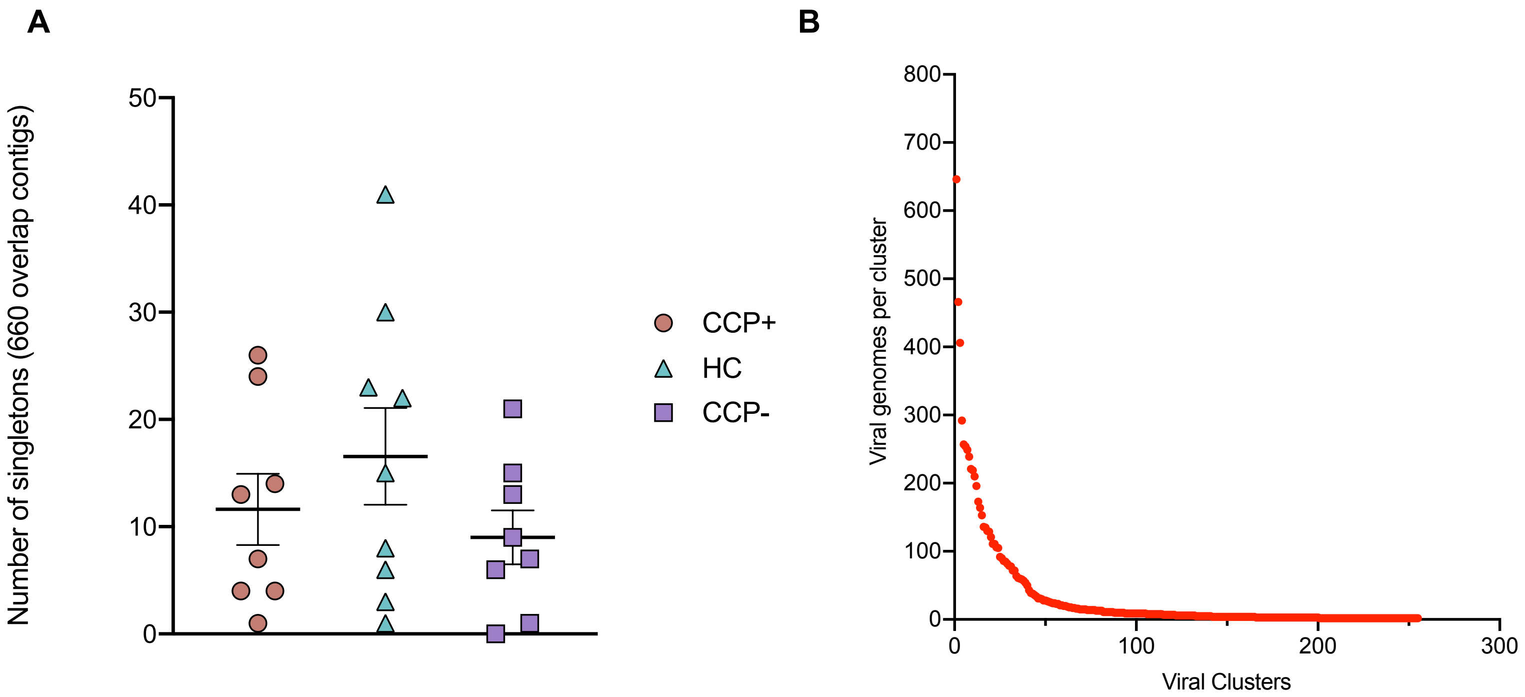

### Supplemental Figure 5

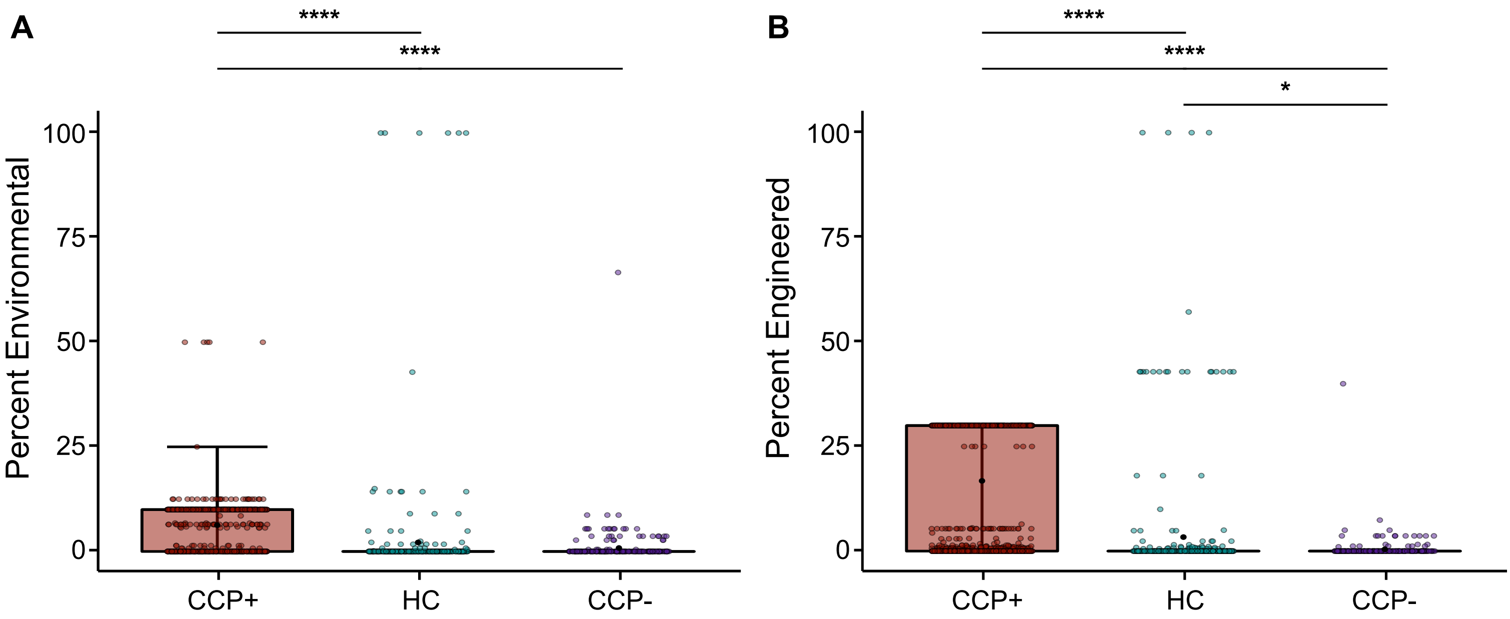

### Supplemental Figure 6

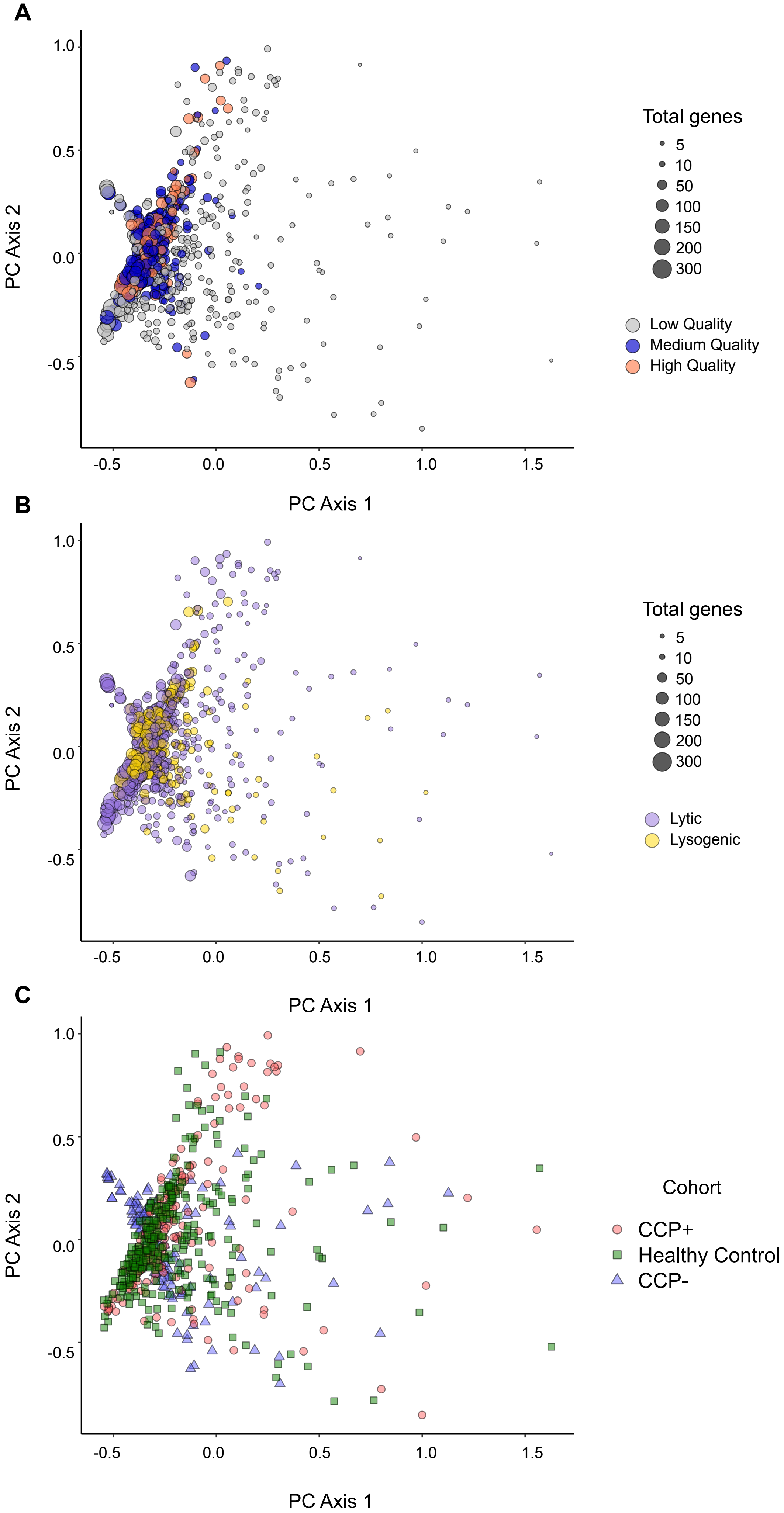

### Supplemental Figure 7

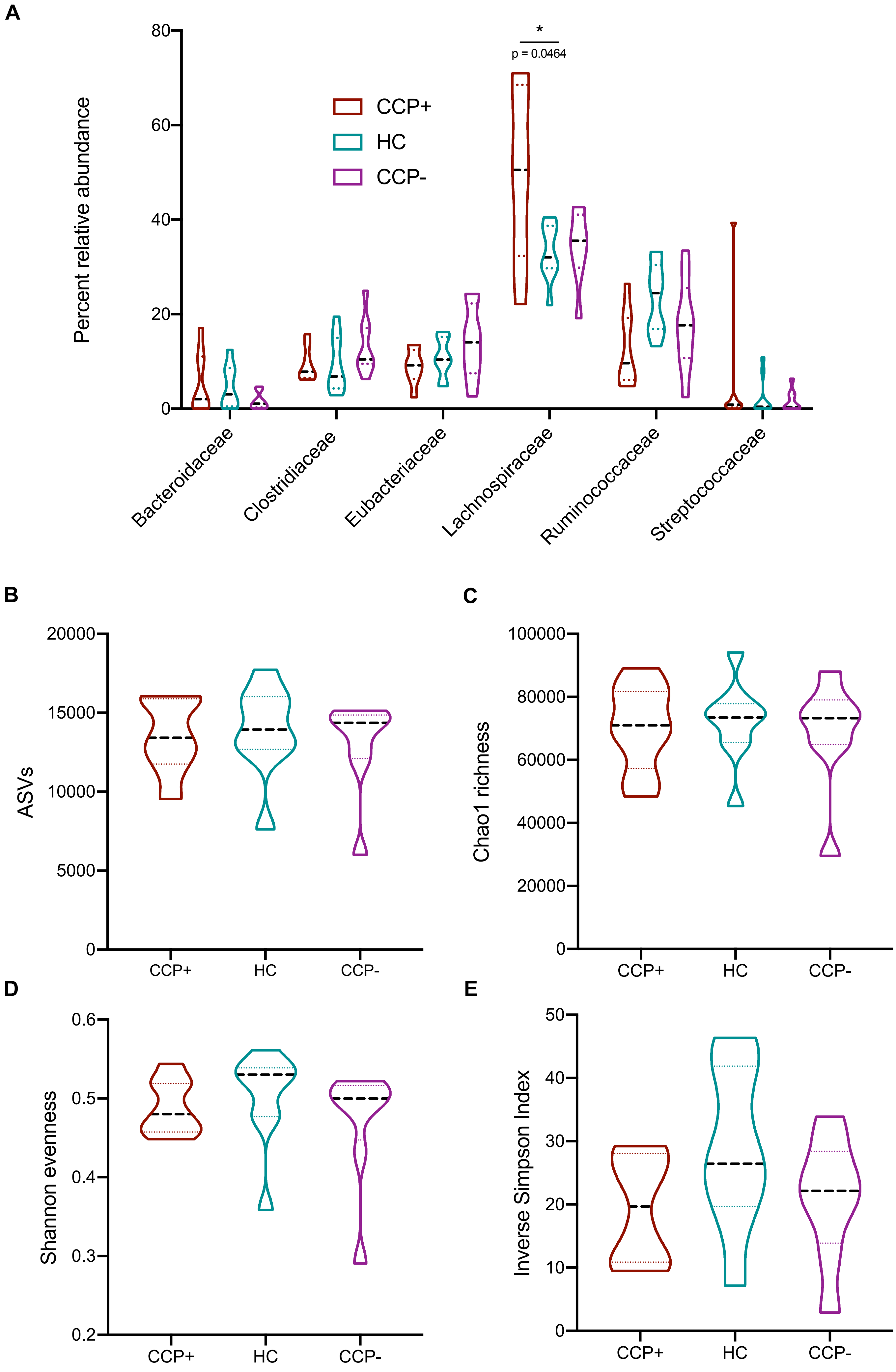

### Supplemental Figure 8

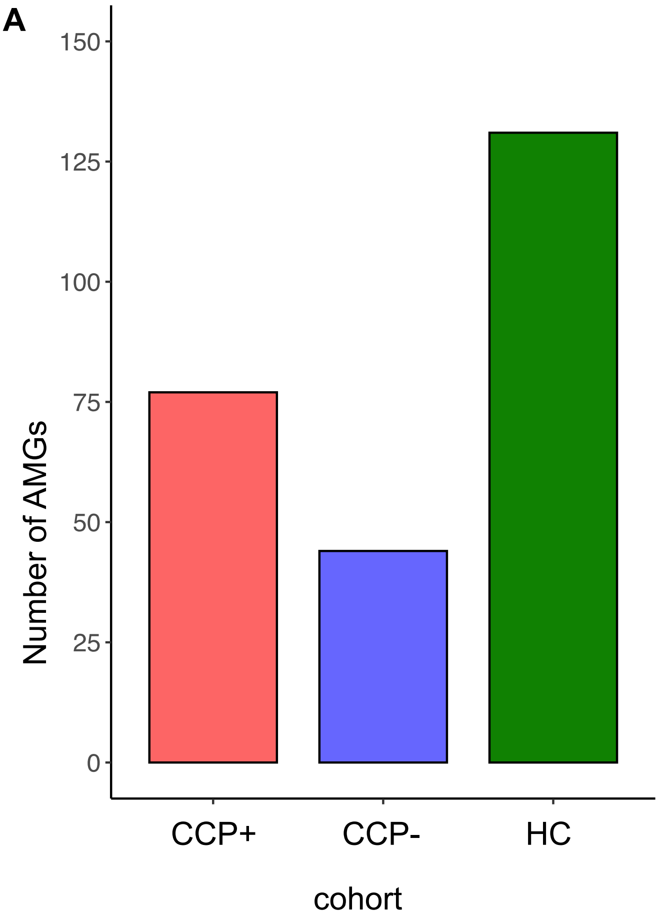

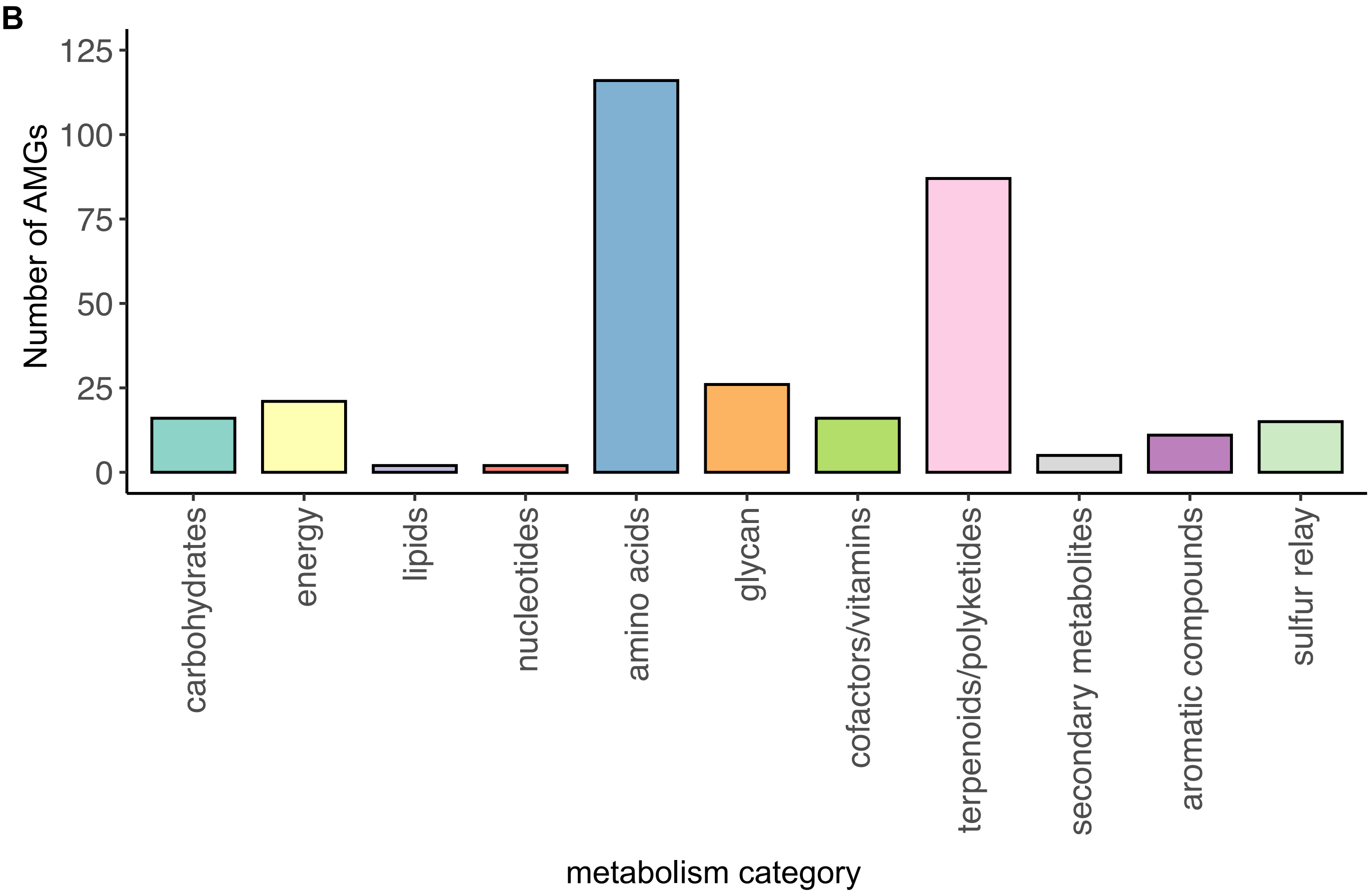
